# *H2A.X* mutants exhibit enhanced DNA demethylation in *Arabidopsis thaliana*

**DOI:** 10.1101/2023.01.08.523178

**Authors:** Jennifer M. Frost, Jaehoon Lee, Ping-Hung Hsieh, Samuel J. H. Lin, Yunsook Min, Matthew Bauer, Anne M. Runkel, Hyung-Taeg Cho, Tzung-Fu Hsieh, Yeonhee Choi, Robert L. Fischer

## Abstract

*H2A.X* is an H2A variant histone in eukaryotes, unique for its ability to respond to DNA damage, initiating the DNA repair pathway. *H2A.X* replacement within the histone octamer is mediated by the FAcilitates Chromatin Transactions (FACT) complex, a key chromatin remodeler. FACT is required for DEMETER (DME)-mediated DNA demethylation at certain loci in *Arabidopsis thaliana* female gametophytes during reproduction, though it is not known how FACT targets DME sites. Here, we investigated whether H2AX is involved in DME- and FACT-mediated DNA demethylation during Arabidopsis reproduction. We show that *h2a.x* mutants are more sensitive to genotoxic stress, consistent with previous reports. *H2A.X* fused to the *Green Fluorescent Protein (GFP)* gene under the *H2A.X* promoter was highly expressed in newly developing *Arabidopsis* tissues, including in male and female gametophytes. We examined DNA methylation in *h2a.x* developing seeds using whole genome bisulfite sequencing, and found that CG DNA methylation in the developing endosperm, but not the embryo, is decreased genome-wide in *h2a.x* mutants, predominately in transposons and intergenic DNA. Hypomethylated sites overlapped 62 % with canonical DME loci. These data indicate that H2A.X is not required for DME function, but is important for DNA methylation homeostasis in the unique chromatin environment of *Arabidopsis* endosperm.

## Introduction

DNA methylation regulates gene expression and silences transposable elements (TEs) in plants and vertebrates (Law and Jacobsen, 2010), and epigenetic reprogramming by DNA demethylation is vital for reproduction in mammals and flowering plants (Monk et al., 1987;Feng et al., 2010;Wu and Zhang, 2017;Parrilla-Doblas et al., 2019). In *Arabidopsis thaliana*, DNA demethylation during reproduction is catalyzed by the DNA glycosylase DEMETER (DME) (Choi et al., 2002). DME is a dual function glycosylase/AP lyase, which actively removes DNA methylation via the Base Excision Repair (BER) pathway (Gehring et al., 2006).

DME-mediated DNA demethylation occurs genome-wide at discrete loci that fall into two groups. The first consists of relatively euchromatic, AT-rich, small TEs that are nucleosome-poor, and generally interspersed with genes in chromosome arms (Ibarra et al., 2012). The second group of loci requires the Facilitates Chromatin Transactions (FACT) complex for DME access, and are longer, heterochromatic TEs prevalent in pericentromeric, gene poor regions, enriched with heterochromatic histone marks and H1 linker proteins (Frost et al., 2018). During reproduction DME and DME-FACT mediated DNA demethylation occurs specifically in male and female gamete companion cells, the vegetative and central cells, respectively (Ibarra et al., 2012;Park et al., 2017), and is vital for *Arabidopsis* reproduction, whereby loss of maternal DME or FACT results in development abnormalities, loss of genomic imprinting and seed abortion (Choi et al., 2002;Gehring et al., 2006;Hsieh et al., 2009;Ikeda et al., 2011;Ibarra et al., 2012).

FACT is required for several other vital cellular functions, including transcription initiation and elongation, nucleosome assembly and disassembly, and for histone variant exchange, specifically of H2A.X (Belotserkovskaya et al., 2003;Heo et al., 2008;Formosa, 2012;Piquet et al., 2018). In *Arabidopsis*, H2A.X is essential for the response to DNA damage, whereby the phosphorylation of its SQEF motif by Ataxia Telangiectasia Mutated (ATM) and ATR kinases, serves as a signal for recruitment of DNA repair and checkpoint proteins (Du et al., 2006;Heo et al., 2008;Dantuma and van Attikum, 2016). It is not known how FACT is recruited to DME target sites, and the apurinic/apyrimidinic (AP) sites created during base-excision repair (BER) can lead to the formation of double strand breaks (Sczepanski et al., 2010). We therefore sought to explore whether recruitment of H2A.X to sites of DME activity during BER may provide a functional link between H2A.X, FACT and DME during *Arabidopsis* reproduction. In order to investigate this, we analyzed the expression and activity of *H2A.X* during *Arabidopsis* reproduction, finding that *H2A.X* is expressed throughout the plant, particularly in developing tissues and the male and female gametophytes. The loss of H2A.X does not impair DME-mediated DNA demethylation, instead leading to CG hypomethylation at DME sites, as well as other intergenic regions and transposable elements, specifically in the endosperm.

## Materials and Methods

### Plant materials and growth conditions

Arabidopsis seeds were bleached and sown onto Murashige and Skoog plates, followed by vernalisation in the dark at 4 degrees for 3 days, and two weeks growth in a light chamber, before transplantation onto soil. Seedlings were grown in a greenhouse with a long-day photoperiod (16 h light, 8 h dark). Seed stocks of T-DNA insertion mutants (SALK_012255 in HTA3 and SAIL_382_B11/CS873648 in HTA5, Figure 1A) in the Columbia-0 (Col-0) background were obtained from the ABRC stock center. Mutant alleles were, backcrossed five times to wild-type, and finally crossed to obtain double *hta3/hta3; hta5/hta5* null plants, designated *h2a.x*, as well as segregating wild-type siblings. Both T-DNA insertion alleles have been studied and validated in recent reports (Lorkovic et al., 2017;Waterworth et al., 2019).

**Figure 1.**
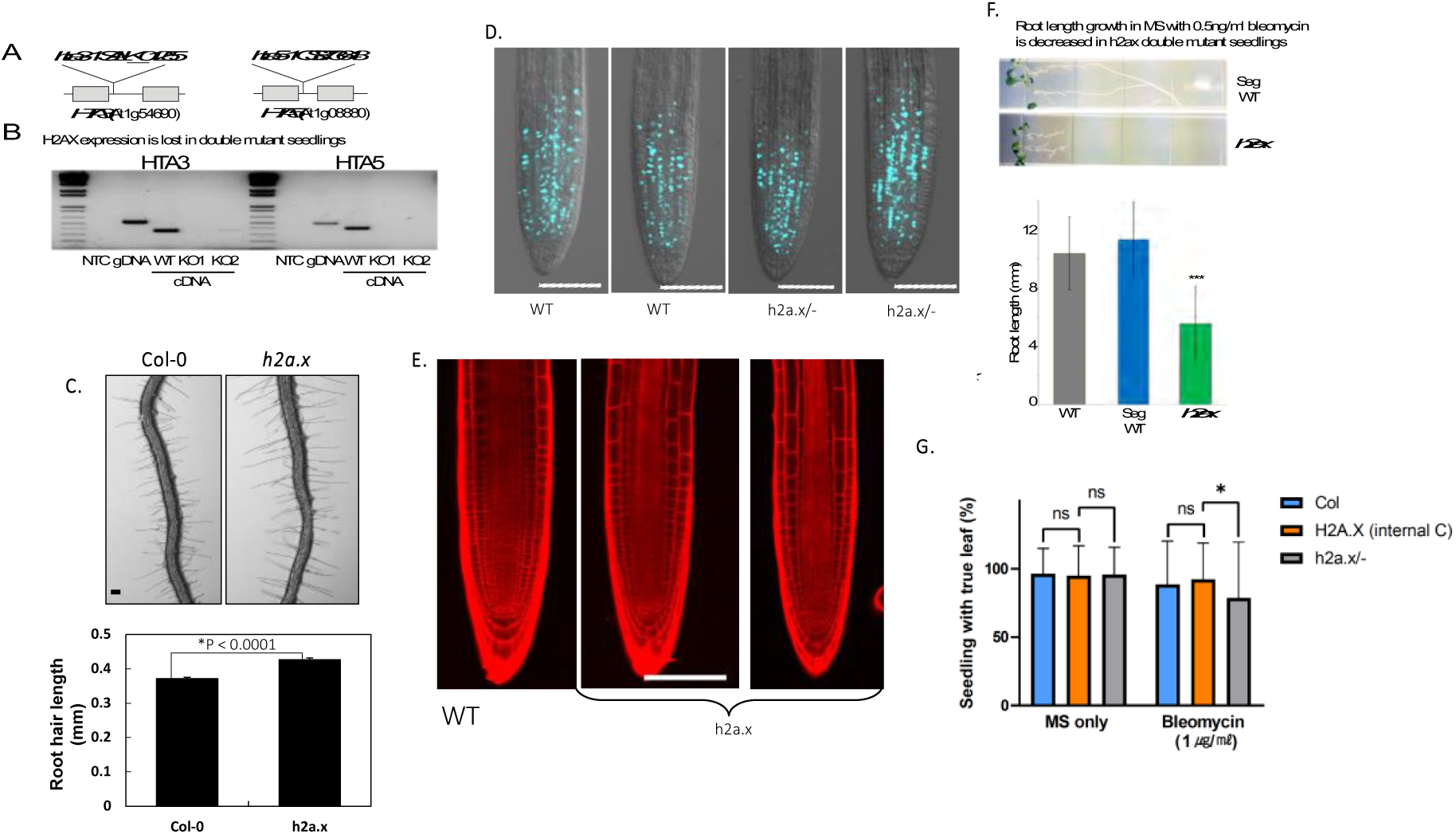
*h2a.x* mutant phenotype analysis. (A)HTA.3 and HTA.5 genomic loci, showing gene structure and location of T-DNA insertions. (B) qPCR analysis of each mutant, showing cDNA-specific PCR amplification and loss of gene product in mutant seedling tissue (C) Root hair phenotypes of wild-type (Col-0) and h2a.x mutant primary roots and length measurements in mm. Data represent mean ± SEM (n = 1,419 root hairs for Col-0 and 1,159 root hairs for h2a.x from 35 ∼ 40 roots. The asterisk (*) indicates a significant difference (Student’s t test). Scale bar, 100 μm. (D) EdU staining of WT and h2a.x double mutant roots at 3 DAG. Scale bar, 100 μm. (E) Propidium iodide (PI) staining of WT and h2a.x double mutant roots. Scale bar, 100 μm. (F) Aberrant root growth of h2a.x mutant seedlings when grown in bleomycin MS. Root length measurements are in mm and the result of three replicate experiments, each with 15 seedlings. (G) The formation of true leaves was slightly reduced in h2a.x mutant seedlings when grown in bleomycin MS. Measurements are the result of duplicated experiments. Leaves counted are; 415 in MS only and 203 in bleomycin MS for Col-0, 395 in MS only and 209 in bleomycin MS for H2A.X internal segregated WT control, 406 in MS only and 202 in bleomycin MS for h2a.xmutant. The box height and whisker length indicate the mean and standard deviation of each sample, respectively. The significance of differences between samples was measured by the Kolmogorov-Smirnov test. ns, not significant; * p=0.0440

### Edu cell proliferation assay

5-ethynyl-2’-deoxyuridine (EdU) staining using an Invitrogen Click-iT™ EdU Alexa Fluor™ 488 HCS Assay (C10350) was performed to detect S phase cells. Seeds were grown in MS media vertically for 3 days. Seedlings were collected in MS solution containing 1μM Edu and incubated at 22°C for 30 minutes. And then, samples were fixed in 4%(w/v) formaldehyde solution in phosphate-buffered saline (PBS) with 0.1% Triton X-100 for 30min, and washed three times with PBS each for 5 minutes. The samples were incubated in Edu detection cocktail solution at room temperature for 30 minutes in the dark, and washed with the Click-iT® rinse buffer and then three times with PBS. The photos were taken using confocal microscopy (LSM700, Zeiss).

### Propidium Iodide (PI) staining

Propidium Iodide (P-4170, sigma) staining was used to detect cell death and show the anatomy of the roots. The samples were stained with working PI solution (5ml PI solution in 1ml of distilled water) at room temperature for 30s and washed with distilled water on slide glass.

### Observation of root hair phenotypes

Root hair phenotypes were observed under a stereomicroscope (M205 FA, Leica). Root hair length was measured as previously described by (Lee and Cho, 2006) with slight modifications. DAG3 seedling roots were photographed digitally using the stereomicroscope at 40X magnification. The hair length of nine consecutive hairs which protruded perpendicularly from each side of the root, for a total of 18 hairs from both sides of the root, was calculated using ImageJ 1.50b software (National Institutes of Health).

### *H2A.X* expression localization

HTA3 and HTA5 GFP fuson proteins were cloned alongside a hygromycin resistance casette using a Gibson assay (Invitrogen) and F1 seeds screened on MS containing hygromycin. F1 plants were screened manually using a fluorescence microscope and seeds collected from plants expressing GFP. F2 seeds were grown on hygromycin and selected if we identified segregation of the resistance allele, indicating the presence of a single copy transgene. F3 plants were then used for confocal microscopy.

### DNA damage assay

*Arabidopsis* seeds were planted on MS containing 0.5ug/ml bleomycin sulphate and grown vertically for 14 days under long day conditions, before measuring root length. MS without bleomycin was used as a control. Values are from three independent experiments each including 15 seedlings for each genotype. Truf leaf assay was performed as previously described with 10-day-old seedlings (Min et al., 2019).

### Isolation of *Arabidopsis* endosperm and embryos

WT Col-0 and *h2a.x* mutant *Arabidopsis* flower buds were emasculated at flower stage 12-13 using fine forceps and pollinated with L*er* pollen 48 hours later. Eight to ten days after pollination (DAP) developing F1 seeds (linear to bending cotyledon stage) were immersed in dissection solution (filter-sterilized 0.3 M sorbitol and 5 mM pH 5.7 MES) on sticky tape and dissected by hand under a stereo-microscope using fine forceps (Fine Science Tools, Inox Dumont #5) and insect mounting pins. The seed coat was discarded, and debris removed by washing collected embryos or endosperm five to six times with dissection solution under the microscope.

### Bisulfite sequencing library construction

As described previously, genomic DNA was isolated from endosperm and embryo (Hsieh et al., 2009). Bisulfite sequencing libraries for Illumina sequencing were constructed as in (Ibarra et al., 2012) with minor modifications. In brief, about 50 ng of genomic DNA was fragmented by sonication, end repaired and ligated to custom-synthesized methylated adapters (Eurofins MWG Operon) according to the manufacturer’s instructions for gDNA library construction (Illumina). Adaptor-ligated libraries were subjected to two successive treatments of sodium bisulfite conversion using the EpiTect Bisulfite kit (Qiagen) as outlined in the manufacturer’s instructions. The bisulfite-converted library was split between two 50 ul reactions and PCR amplified using the following conditions: 2.5 U of ExTaq DNA polymerase (Takara Bio), 5 μl of 10X Extaq reaction buffer, 25 μM dNTPs, 1 μl Primer 1.1 and 1 μl multiplexed indexing primer. PCR reactions were carried out as follows: 95ºC for 3 minutes, then 14-16 cycles of 95 ºC 30 s, 65 ºC 30 s and 72 ºC 60 s. Enriched libraries were purified twice with AMPure beads (Beckman Coulter) prior to quantification with the Qubit fluorometer (Thermo Scientific) and quality assessment using the DNA Bioanalyzer high sensitivity DNA assay (Agilent). Sequencing on either the Illumina HiSeq 2000/2500 or HiSeq 4000 platforms was performed at the Vincent J. Coates Genomic Sequencing Laboratory at UC Berkeley.

### Bisulfite data analysis

Sequenced reads were sorted and mapped to the TAIR8 Col-0 and Ler genomes in cases of seeds derived from Col x Ler crosses, or not sorted and mapped to TAIR8 Col-0. Gene and TE ends analysis and kernel density plots were generated as previously described (Ibarra et al., 2012), using only windows with at least 10 informative sequenced cytosines, and fractional methylation of at least 0.7 (CG), 0.4 (CHG) or 0.08 (CHH) in at least one of the samples being compared.

## Results

### *Arabidopsis* seedlings lacking *H2A.X* have reduced DNA damage tolerance

*H2A.X* is encoded by two genes in *Arabidopsis*, HTA3 (AT1G54690) and HTA5 (AT1G08880). To investigate the effect of *H2A.X* mutations, we generated double mutants lacking both HTA3 and HTA5, verified the loss of transcripts using RT-PCR (Figure 1B) and analyzed the sporophytic and gametophytic phenotypes of *h2a.x* plants. *h2a.x* mutant allele segregation, plant morphology, growth rate and flowering time were all normal, except for a significant increase in root hair length (Figure 1C, *p*<0.0001). *h2a.x* root hairs were ∼15 % longer than WT three days after germination (DAG). We then tested whether cell proliferation was normal in *h2a.x*, using the 5-ethynyl-2′-deoxyuridine (EdU), a thymine analog, and click chemistry to measure incorporation in newly synthesized DNA (Kotogany et al., 2010). We did not observe a difference in EdU-stained cells between wild-type and *h2a.x* roots (Figure 1D), indicating that S-phase progression and cell proliferation are normal in *h2a.x* mutants. We also measured whether there was increased DNA damage in mutant roots using propidium iodide (PI) staining but did not observe any differences between WT and *h2a.x* (Figure 1E). These observations are consistent with mutant phenotypes observed in other DNA damage pathway genes, such as ATM or ATR kinases, which only exhibit a phenotype under growth conditions that promote DNA damage (Culligan et al., 2006).

We therefore grew *h2a.x* and segregating WT seeds on MS plates containing Bleomycin sulphate, which induces double strand breaks (DSB) in DNA. MS Bleomycin concentrations of 0.5 ng/ml were used to test primary root formation and 1 ug/ml to test true leaf formation, as root development was more sensitive to the drug. *h2a.x* mutant seedlings had a significant reduction in root length compared to WT (Figure 1F). True leaf formation rate is slightly reduced in *h2a.x* mutants (Figure 1G). These data are consistent with previous findings, also showing aberrant true leaf and root growth in *h2a.x* double mutant seedlings grown under genotoxic stress (Lorkovic et al., 2017). Thus, a lack of *H2A.X* resulted in increased sensitivity of developing tissues to DNA damaging agents, showing that *H2A.X* is required for the response to DNA damage in *Arabidopsis*.

### *H2A.X* is widely expressed across *Arabidopsis* tissues, including in gamete companion cells

To investigate the role of *H2A.X* in *Arabidopsis* development, we analyzed its expression pattern in sporophytic and reproductive tissues. We generated translational fusion constructs between the Green Fluorescent Protein (GFP) gene and the HTA3 and HTA5 genes, including their promoter sequences, and introduced them into WT Col-0 *Arabidopsis* plants using Agrobacterium mediated transfer, deriving three and four independent lines for each allele, respectively. GFP fluorescence was observed using confocal microscopy. Both HTA3 and HTA5 were expressed in dividing cells of the sporophyte: First true leaves (Figure *2*A and B), the floral meristem (Figure 2C), the adaxial leaf surface (Figure 2D and E), petal tips (Figure 2F), in root meristem and root tips (Figure 2G and H). In reproductive structures supporting gametophyte development, such as the ovule primordia (Figure 2I), ovules (Figure 2J), and anthers (Figure 2K) both isoforms were expressed. In the next generation seed, both isoforms were present in the developing embryo, but not in the endosperm (Figure 2L and M). In each case, HTA5 expression was more prevalent and more widely expressed than HTA3. Conversely, in gametophytic development, both isoforms were again expressed but HTA3 was the dominant isoform (Figure 3). In the male gametophyte, both HTA3 and HTA5 were present in the microspore prior to mitosis. After Pollen Mitosis 1 (PMI) HTA3 was expressed in the generative and vegetative nucleus of bicellular pollen, and following Pollen Mitosis II (PMII), in both sperm cells and the vegetative nucleus of mature, tricellular pollen (Figure 3A). HTA5 expression was also present in both the generative and vegetative nucleus following PMI, but was lost in the vegetative nucleus following PMII, in tricellular pollen (Figure 3B). In the female gametophyte, egg cell expression was visible for both HTA3 and 5, but was weak, conversely, HTA3 expression was very striking in the central cell, where it persisted following fertilization in the first cell divisions of the developing endosperm (Figure 3C and E). HTA5 expression was also observed in the central cell, but expression in the surrounding ovule tissue was more prominent for this isoform (Figure 3D).

**Figure 2.**
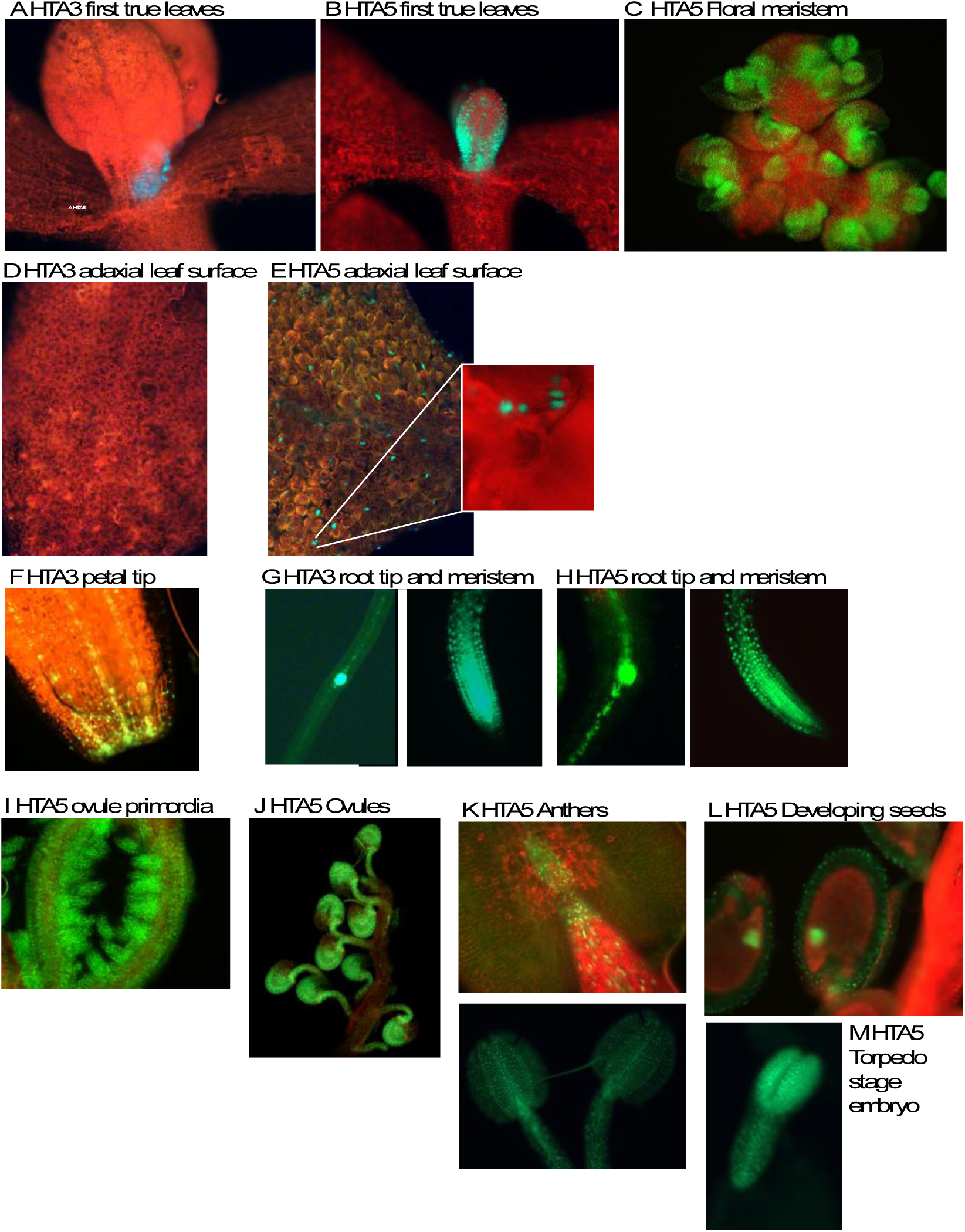
*HTA5* expression is more dominant in dividing cells of the sporophyte. First true leaves (A and B), the floral meristem (C), the adaxial leaf surface (D and E), petal tips (F), in root meristem and root tips (G and H). In reproductive structures supporting gametophyte development such as the ovule primordia (I), ovules (J), and anthers (K), both isoforms were expressed. In the next generation seeds, both isoforms were present in the developing embryos (L and M).

**Figure 3.**
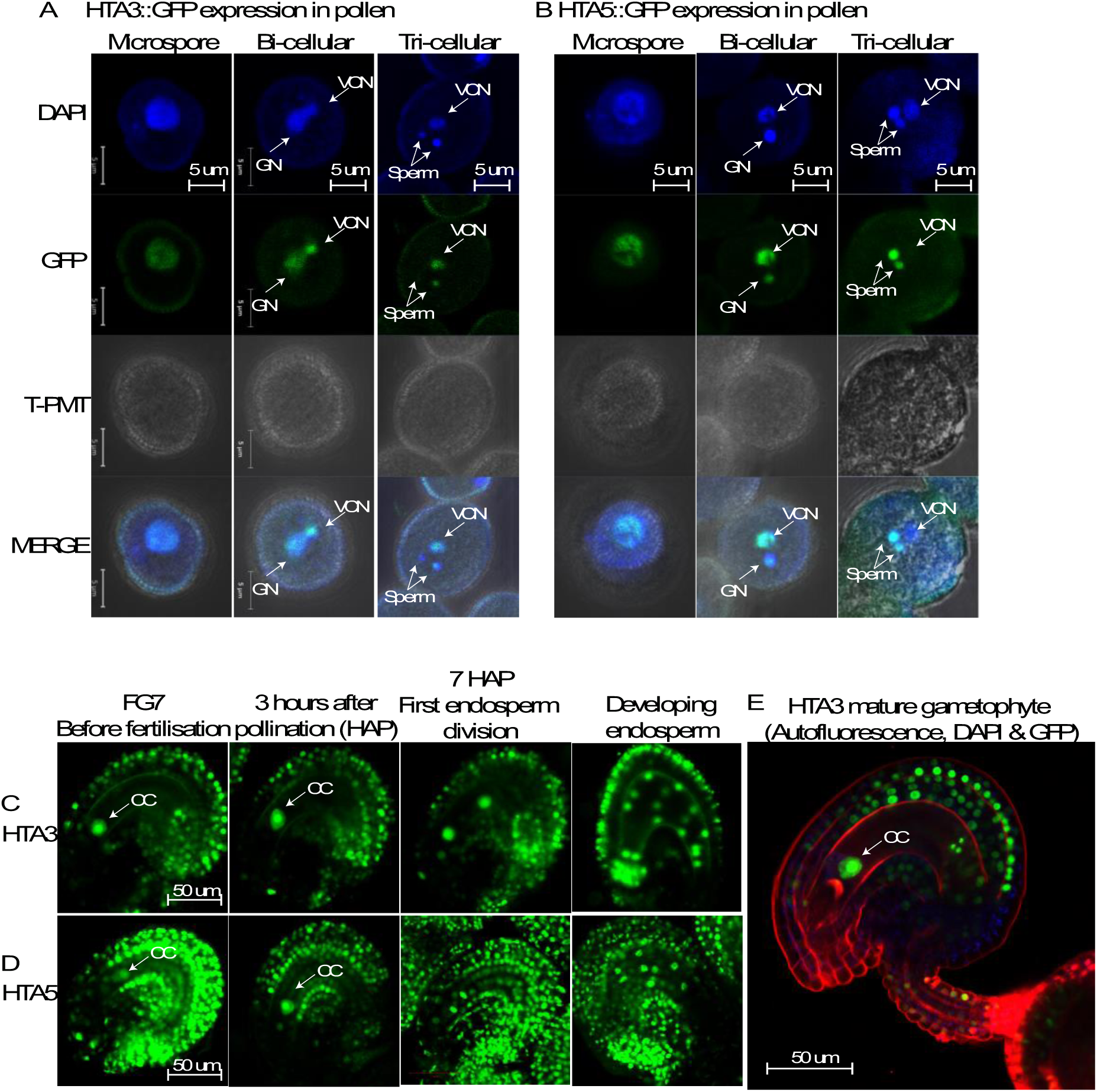
*HTA3* expression is more dominant in gametophytic development. In the male gametophyte, both HTA3 and HTA5 are present in the microspore prior to mitosis. After Pollen Mitosis I (PMI) *HTA3* was expressed in the generative and vegetative nucleus of bicellular pollen, and following Pollen Mitosis II (PMII), in both sperm cells and the vegetative nucleus of mature, tricellular pollen (A). *HTA5* expression was also present in both the generative and vegetative nucleus following PMI, but was lost in the vegetative nucleus following PMII, in tricellular pollen (B). In the female gametophyte, egg cell expression was visible for both *HTA3* and *HTA5*, but was weak. Conversely, *HTA3* expression was very striking in the central cell, where it persisted following fertilization in the first cell divisions of the developing endosperm (C and E). *HTA5* expression was also observed in the central cell, but expression in the surrounding ovule tissue was more prominent for this isoform (D).

### DME does not regulate *H2A.X* expression in the Arabidopsis gametophyte

Since *H2A.X* expression was prominent in the central and vegetative cells, specifically where DME-mediated demethylation and related BER activity takes place (Schoft et al., 2011;Park et al., 2017), we reasoned that *H2A.X* expression may be regulated by promoter DNA methylation, whereby DME might demethylate *HTA3* and *HTA5* promoter sequences in the gametophyte, increasing expression of these transcripts. To test this hypothesis, we utilized wild-type plants hemizygous for the *HTA3:GFP* transgene, for which strong HTA3 expression could be observed in the central cell in ∼50 % of the developing ovules (Figure 4). We crossed these plants with DME/*dme*-2 heterozygotes to derive DME/*dme*-2 plants that were also hemizygous for the *HTA3:GFP* transgene. A maternally inherited *dme*-2 mutation generates embryo abortion and seed lethality, so analysis of seed development is generally only possible in DME/*dme-2* heterozygotes. We then analyzed the incidence of HTA3:GFP expression in DME/*dme-2* mutants and their segregating wild-type siblings in the F2 population. In both DME/DME HTA3:GFP/- (Figure 4A) and DME/*dme*-2 HTA3:GFP/- F2 (Figure 4B) siblings we observed that ∼50 % of the female gametophytes within ovules produced a strong GFP signal, indicating that the loss of DME did not alter the expression of *H2A.X* in the *Arabidopsis* female gametophyte. Consistent with this, when we compared promoter DNA methylation for the H2A.X variants in *Arabidopsis* wild-type and *dme-2* mutant central cells and endosperm (Hsieh et al., 2009;Park et al., 2016), we found that *H2A.X* promoter methylation was low in both tissues, and unchanged in the *dme*-2 mutant (Supplementary Figure S1A and 1B). Other H2A variant gene loci were also unmethylated in both wild-type and *dme-2* mutant central cell and endosperm, except for H2A.Z.4, which exhibited promoter methylation in central cell and endosperm, that increased in *dme-2* mutants, a hallmark of a DME-target promoter (Supplementary Figure S1C).

**Figure 4.**
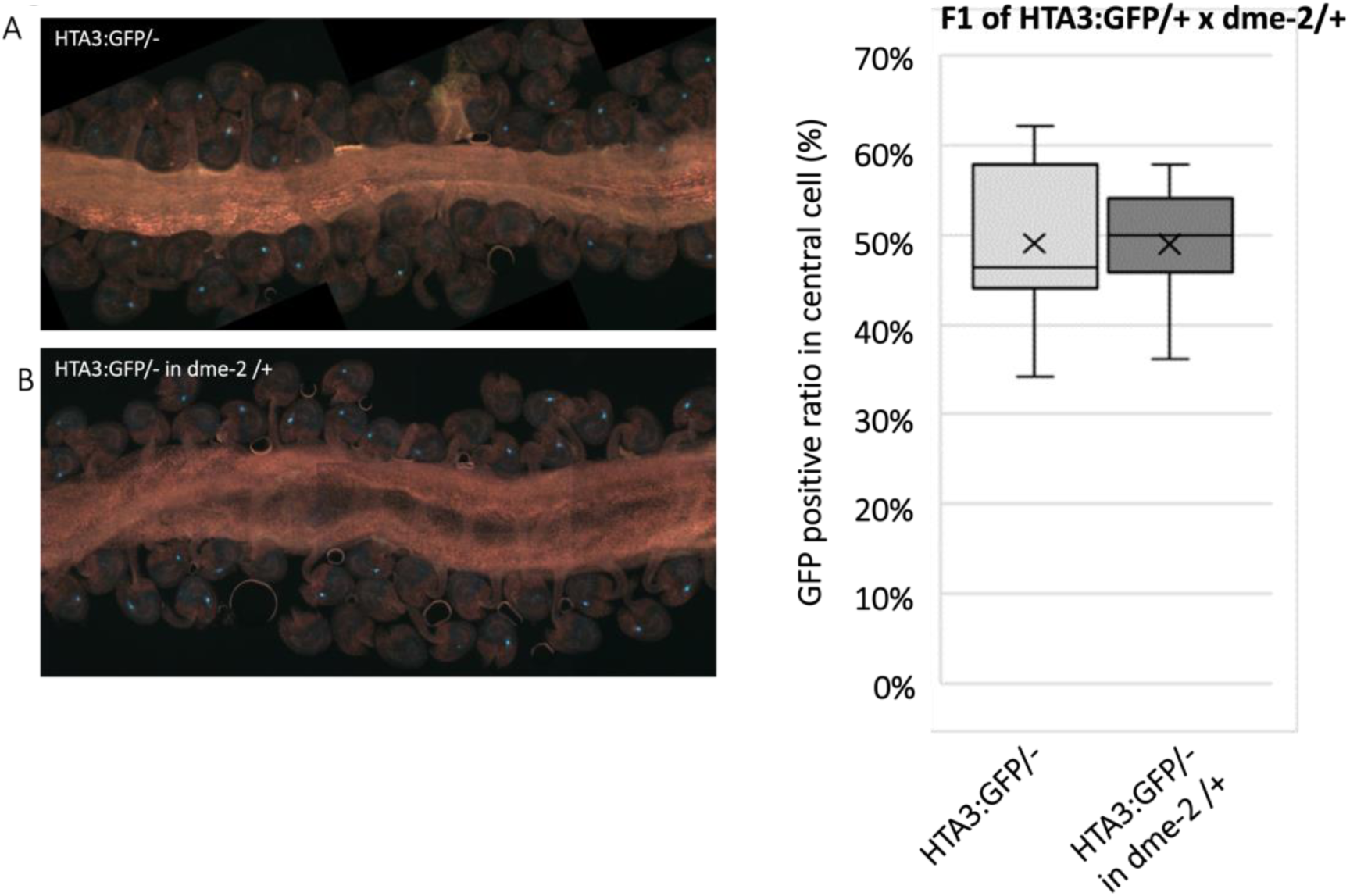
*HTA.3:GFP* expression in WT and *dme-2* mutant central cells. (A) Confocal image of WT and *dme-2* mutant developing ovules expressing an *HTA.3:GFP* transgene. Expression is confined to the central cell. (B) Box plot showing the distribution of GFP positive central cells between WT and *dme-2* mutant ovules. There was no significant change in GFP positive central cells in DME/*dme-2*. + mark in boxplot is mean of data. The line in a box is median (n = 554 mature ovules for HTA3:GFP/- plants and 412 mature ovules for HTA3:GFP/- in DME/*dme-2* heterozygous plants. The values are not significantly different by Student’s *t* test.

### *h2a.x* mutant endosperm is hypomethylated genome-wide

To investigate whether changes in DNA methylation were present in *h2a.x*, we carried out genome-wide bisulfite sequencing (BS-seq) of manually dissected endosperm and embryo from homozygous *h2a.x* mutant and wild-type F1 seeds and their resulting seedlings, following self-pollination of homozygous *h2a.x* mutants and wild-type sibling plants. We observed that embryo DNA methylation in the *h2a.x* mutant was unchanged from wild-type, with the peak of fractional methylation difference at zero (Figure 5A). However, DNA methylation of *h2a.x* mutant endosperm was reduced compared to wild-type in the CG context, with the fractional methylation difference peak shifted to the left (Figure 5B).

**Figure 5.**
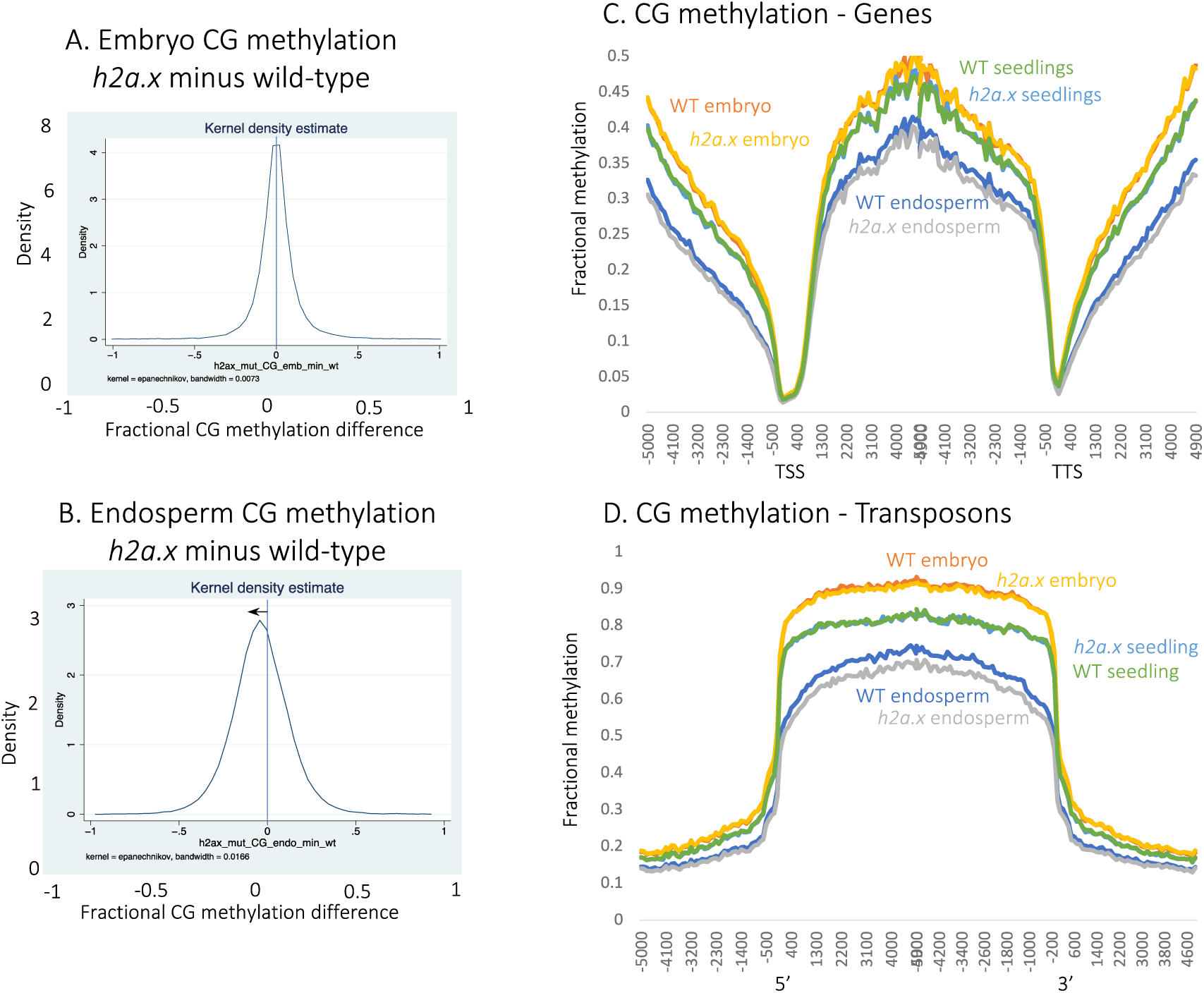
Genome-wide methylation analysis of selfed double *h2a.x* mutant developing embryo, endosperm and seedling. (A) Fractional methylation difference between *h2a.x* double mutant and WT CG methylation from embryo (linear-bending cotyledon) is plotted, data in 50 bp windows with >10x sequence coverage. Data are from *h2a.x* Col selfed plants and segregating wild type siblings. (B) As for A, but with endosperm. (C) Ends analysis of *h2a.x* mutant genomic methylation in genes, with genes aligned according to their 5’ and 3’ ends. (D) Ends analysis of *h2a.x* mutant genomic methylation in transposons, with transposons aligned according to their 5’ and 3’ ends.

To ascertain which endosperm loci were hypomethylated, we aligned our methylome data to the 5’ transcriptional start sites (TSS) and 3’ transcriptional termination sites (TTS) of genes and transposons, and also included methylome of *h2a.x* seedlings, which also showed that hypomethylation was present only in endosperm, and although present in gene bodies and intergenic regions, was most striking in transposon bodies, (Figure 5C and D). CHG and CHH methylation was also reduced in endosperm transposon bodies (Supplementary Figure S2A-D). In *h2a.x* embryos, CHH methylation in TEs was decreased (Figure S5D), although embryo CHH methylation increases steadily with time during embryo development (Papareddy et al., 2020) so it is possible the differences observed in embryos are a technical difference, whereby mutant seeds were dissected earlier than wild-type.

### H2A.X hypomethylation overlaps DME target loci

Inheritance of a loss-of-function maternal *dme* allele or a loss-of-function maternal *ssrp1* allele, which encodes one of the proteins in the FACT complex, results in a striking phenotype of seed abortion and developmental delay. Seed viability in homozygous *h2a.x* mutants, as well as in crosses from maternal *h2a.x* with wild-type Col-0 pollen, was normal, suggesting that DME- and DME-FACT-mediated DNA demethylation occurred normally in *h2a.x* mutant seeds, at least in the DME-regulated PRC2 genes critical for seed viability. In wild type female gametophytes, DME and DME-FACT mediated DNA demethylation leads to a hypomethylated endosperm compared to embryo (Hsieh et al., 2009;Ibarra et al., 2012;Park et al., 2016;Frost et al., 2018). To assess whether *h2a.x* hypomethylation may still be related to DME activity, we compared differential methylated regions (DMRs) between endosperm and embryo in WT and *h2a.x* mutant seeds. There were 4,451 hypo-DMRs between WT endosperm vs embryo, covering about 1.3 M bps, which reflect the activity of DME in the central cell. In contrast, 7,526 hypo-DMRs were identified between *h2a.x* endosperm vs embryo, covering 2.7 M bps in length, more than double the area of the wild-type hypo-DMRs (Figure 6A). Of these, 4692 (62 %) overlapped with canonical DME DMR loci (Ibarra et al., 2012). The hypo-DMRs consisted of both WT embryo-endosperm DMRs (n=3238) and novel *h2a.x* specific DMRs (n = 4357, Figure 7A and B). There was also a group of WT DMRs, which were only differentially methylated between WT embryo and endosperm (n=1213). We delineated DMRs by size (0.1 kb->1.5 kb) and found that *h2a.x* embryo-endosperm DMRs were represented across all size classes (Figure 6C).

**Figure 6.**
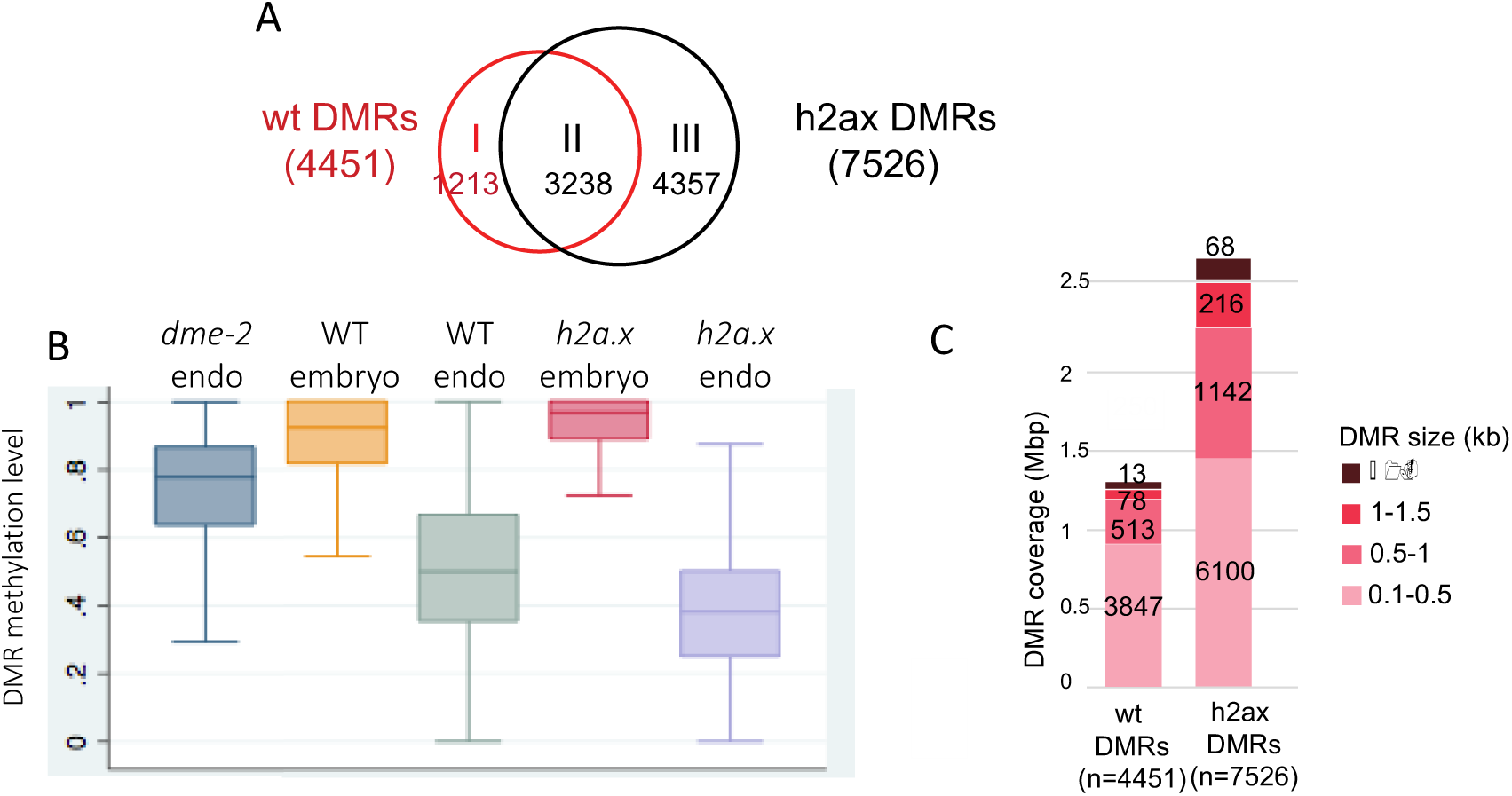
Endosperm-Embryo hypomethylated DMR analysis in WT and h2ax mutants. Analysis of h2a.x methylomes, comparing DMRs between endosperm and embryo. (A) Venn diagram illustrating that WT embryo and endosperm harbour 4451 DMRs, the majority of which (3238) are shared with h2a.x embryo and endosperm. h2a.x embryo and endosperm have an additional 4357 DMRs. (B) Box plots showing the relative methylation level of DMRs in embryo and endosperm, in wild type, h2a.x and dme-2 mutants (C) Characterization of h2a.x-specific embryo-endosperm DMRs; wild-type and *h2a.x* Endosperm-Embryo DMRs grouped by size, with the cumulative total length they covered shown, whereby they are represented across all DMR sizes, and represent an overall increase in size distribution.

**Figure 7:**
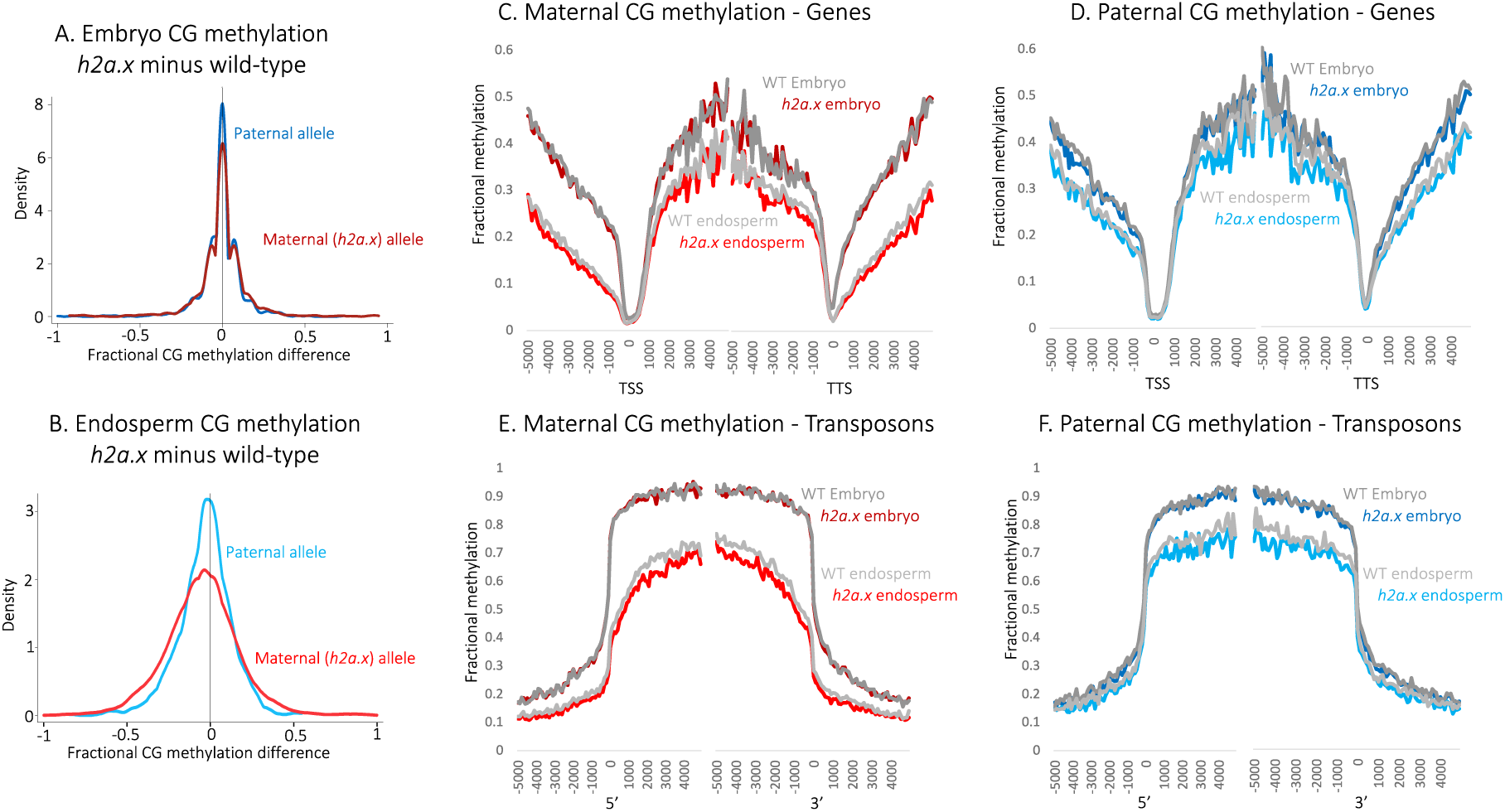
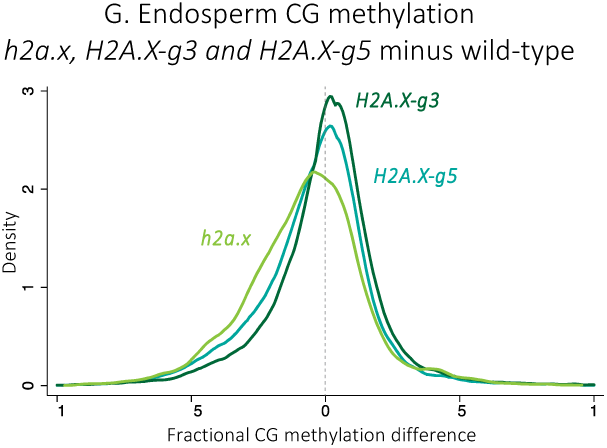
Methylome analysis of seeds *h2a.x* maternal and WT paternal plant crosses. Genome-wide methylation analysis of *h2a.x* mutant developing embryo and endosperm, comparing maternal and paternal alleles in wild-type (WT) and *h2a.x* mutant crosses whereby the maternal allele is either WT Columbia or *h2a.x* homozygous mutant Columbia, and the paternal allele is always wild-type Ler; ‘*h2a.x* paternal’ denotes a wild-type paternal allele now resident in a heterozygous *h2a.x* mutant seed. (A) Fractional methylation difference between *h2a.x* double mutant gametophyte crossed with Ler WT pollen in CG methylation from embryo (linear-bending cotyledon) is plotted, data in 50 bp windows with >10x sequence coverage (B) As for A, but with endosperm. A slight shift towards the left can be seen for the maternal endosperm allele (inherited from *h2a.x* mutant central cell). (C) Ends analysis of maternal (*h2a.x* mutant) genomic methylation in genes, with genes aligned according to their 5’ and 3’ ends. (D) Ends analysis of paternal genomic methylation in genes, with genes aligned according to their 5’ and 3’ ends. (E) Ends analysis of maternal (*h2a.x* mutant) genomic methylation in transposons, with transposons aligned according to their 5’ and 3’ ends. (F) Ends analysis of paternal genomic methylation in transposons, with transposons aligned according to their 5’ and 3’ ends. (G) Fractional CG DNA methylation difference between mutant and WT maternal endosperm in developing seeds from both *hta3/hta3 hta5/+* (*H2A.X-g3*) and *hta5/hta5 hta3/+* mutants (*H2A.X-g5*) were crossed to Ler, so that the sporophyte had one remaining copy of one of the isoforms, but both H2A.X isoforms are lost in ½ of the gametophytes. *H2A.X-g5 and H2A.X-g3* are plotted alongside *h2a.x*. For both *H2A.X-g3 and H2A.X-g5*, the curve peaks are close to zero, whereas the full h2a.x mutant is skewed to the left, indicating redundancy between HTA5 and HTA3 isoforms in the context of mutant endosperm DNA hypomethylation.

### *h2a.x* mutant endosperm hypomethylation is not allele-specific

Seeds are formed following fertilization of both egg and central cell with sperm, resulting in the diploid zygote and triploid endosperm, respectively. Since *h2a.x* hypomethylation is confined to the endosperm, we deduced that this effect may to result from a loss of *h2a.x* in the central cell. Consistent with this idea, both *H2A.X* isoforms are strongly expressed in the wild-type central cell (Figure 3C and D). To identify changes in DNA methylation in the *h2a.x* central cell, we pollinated maternal *h2a.x* mutant plants in the Columbia ecotype with wild-type L*er* pollen. We manually-dissected embryo and endosperm from mutant and segregating wild-type seeds and following BS-Seq, sorted the reads according to their parental ecotype. In this way, the maternal endosperm genome can be used as a proxy for the central cell genome. In *h2a.x* embryos, both maternal and paternal allele CG methylation is identical to WT (peak aligns on zero. Figure 7A), consistent with our observations in self-pollinated *h2a.x* mutants (Figure 5A). However, in endosperm, a slight shift is visible towards the left, indicating mutant hypomethylation, present for both maternal and paternal endosperm (Figure 7B). This indicates that whilst hypomethylation may be inherited from the central cell, resulting in hypomethylated maternal alleles, hypomethylation of the paternal allele must manifest post-fertilization, perhaps due to a reduction in CG methylation efficiency or maintenance. To ascertain which loci were hypomethylated, we again aligned our methylomes to the TSS/5’ and TTS/3’ ends of genes and transposons (Figure 7C-F and Supplementary Figure S3A-D). As in the previous analysis, CG methylation in embryo is not different from wild-type in *h2a.x* mutant gene and transposon bodies, but both maternal and paternal *h2a.x* endosperm alleles are hypomethylated in genes, intergenic regions and TEs, with hypomethylation in TE bodies being most striking. CHG methylation is the same in wild-type and *h2a.x* mutant embryo and endosperm (Supplementary Figure S3E-H) whereas CHH methylation is decreased on both parental alleles, in both *h2a.x* embryo and endosperm (Supplementary Figure S3I-L).

*H2A.X* is encoded by two almost identical isoforms, HTA3 and HTA5.To determine whether one isoform may have an effect independent of the other, we dissected developing seeds from both *hta3/hta3 hta5/+* (*H2A.X-g3*) and *hta3/+ hta5/hta5* mutants (*H2A.X-g5*), crossed to Ler, so that the sporophyte had one remaining copy of one of the isoforms, but both *H2A.X* isoforms are lost in ½ of the gametophytes. Following BS-seq, we determined that both isoforms act redundantly, whereby endosperm methylation was not substantially affected in either *hta3/hta3 hta5/+* or *hta5/hta5 hta3/+* seeds (Figure 7G, H2A.X-g3 and H2A.X-g5, kdensity peaks on zero) compared to hypomethylated *h2a.x* double mutant endosperm (Figure 7G, *h2a.x* peak shifted to the left).

### *h2a.x* hypomethylation is widespread in intergenic DNA

To assess if *h2a.x* endosperm hypomethylated loci are associated with particular chromatin states, we used the published histone marks and genomic characteristics that topologically group the *Arabidopsis* genome into 9 distinct chromatin states (Sequeira-Mendes et al., 2014) and used them to compare methylation differences between *h2a.x* vs wildtype endosperm. For the hypo-DMRs specific to *h2a.x* endosperm vs embryo, the majority reside in non-coding, intergenic sequences, including distal promoters (chromatin state 4, Figure 8A) and AT-rich heterochromatic regions (chromatin states 8 and 9, Figure 8A), consistent with what we observed in TE metaplots (Figure 5C and D, Figure 6E and F). In addition, when we used fractional methylation differences to analyse the chromatin states of hypomethylated loci unique to the *h2a.x* mutant, i.e. not including those present in between wild-type embryo and endosperm, chromatin states 4 and 8 exhibit the largest shift (Figure 8B and C). These data indicate that the novel, *h2a.x*-specific DMRs lie primarily in chromatin states 4, 8 and 9.

**Figure 8:**
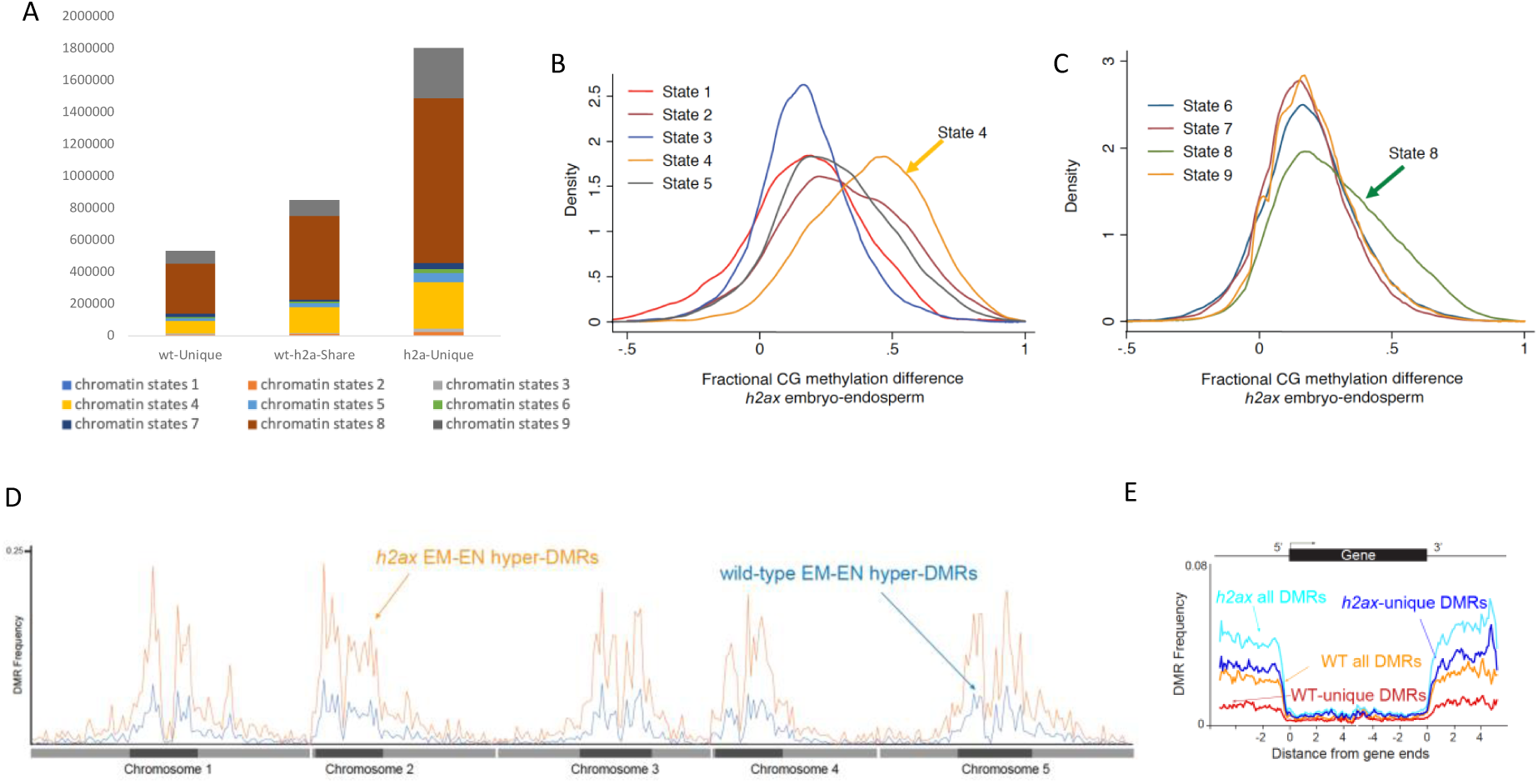
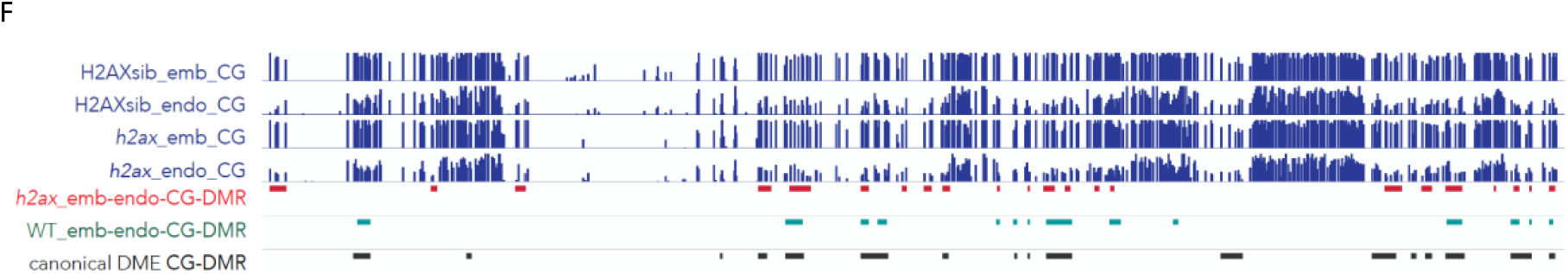
Chromatin states of h2a.x DMRs. (A) Comparison of the chromatin states comprising each group of DMRs from Figure 6A. Chromatin state distribution, and the total length they covered, within wt-unique, wt-*h2a.x* shared, and *h2a.x*-unique Endosperm-Embryo DMRs. States 1 to 7 correspond to euchromatin, and states 8 and 9 correspond to AT- and GC-rich heterochromatin, respectively. The chromatin states most increased (as a fraction of their total) in h2a.x embryo-endosperm DMRs are 4, 8 and 9. (B and C) Kernel density plots showing the fractional methylation difference for h2a.x mutant embryo minus endosperm, plotted according to chromatin state, demonstrating that the largest shift to endosperm hypomethylation (i.e. to the right) lies in State 4 (yellow in B) and State 8 (green in C). (D) Arabidopsis chromosome view of genome-wide methylation levels for h2a.x mutant DMRs between embryo and endosperm, and WT DMRs between embryo and endosperm, represented by the distribution frequency of DMRs along the 5 chromosomes. Dark blocks represent centromere and peri-centromeric regions of each chromosome. (E) Ends analysis plot showing distribution frequency of DMRs with respect to coding genes. Genes were aligned at the 5’- or the 3’-end, and the proportion of genes with DMRs in each 100-bp interval is plotted. DMR distribution is shown with respect to all wt-DMRs (orange trace), wt-unique DMRs (red trace), all *h2a.x*-DMRs (cyan trace), and *h2a.x*-unique DMRs (blue trace). (F) IGV browser view of methylome data and DMR calls for H2AX segregating wild-type embryo and endosperm (Green), h2a.x mutant embryo and endosperm (Red), as well as DMRs identified between dme-2 /wt endosperm (Black) (Ibarra et al.,2012).

In order to characterize the location of *h2a.x* embryo-endosperm DMRs, we plotted their co-ordinates across the *Arabidopsis* genome in 300 kb bins (see Materials and Methods; Figure 8D). This analysis showed that *h2a.x* DMRs, in general, mirror the distribution of wild-type (DME-mediated) embryo-endosperm DMRs, which are enriched in pericentric heterochromatin (Figure 8D). To gain further resolution on DMR location, we aligned wild-type and *h2a.x* DMR coordinates according to 5’ and 3’ ends of genes, which revealed that *h2a.x* hypomethylation is enriched in intergenic regions, consistent with its enrichment in chromatin states 4 and 8 (Figure 8E). To determine whether the *h2a.x* endosperm hypomethylation represented novel sites of demethylation, or resulted from increased demethylation at already demethylated sites (e.g., resulting in longer DMRs), we took a locus-specific approach, using the IGV genome browser to view aligned methylation data and DMRs (Figure 8F). The majority of *h2a.x*-specific hypomethylation represented stand-alone, novel DMRs (red outline). *h2a.x* DMRs overlapped DME-mediated wild-type endosperm/embryo DMRs (green outline), but did not make them longer.

## Discussion

*H2A.X* is one of the H2A variants in higher eukaryotes and differs from canonical H2A by its rapid phosphorylation to y-*H2A.X* in response to DNA double-strand breaks. Unmodified *H2A.X* is ubiquitously expressed and distributed throughout the genome as a component of nucleosome core structure, estimated to represent approximately 10 % of H2A variants present in chromatin at any given time (Rogakou and Sekeri-Pataryas, 1999;Celeste et al., 2003a;Celeste et al., 2003b;Seo et al., 2012). We show that H2AX is widely expressed in *Arabidopsis* newly developing tissues and reproductive cells including the companion cells of the male and female gametophytes. Loss of H2A.X results in endosperm hypomethylation at intergenic regions, and heterochromatic TEs that include DME target sites, revelaing a potential link between H2AX deposition and DME-mediated DNA demethylation.

There are some precedents for H2A variant interactions with DME-like proteins in *Arabidopsis*: H2A.Z is present at the transcriptional start sites of genes, where it promotes transcription by preventing DNA methylation (Zilberman et al., 2008). ROS1/DML2/DML3 act to remove DNA methylation in vegetative tissues and are recruited to a subset of their targets by the IDM1 complex, although they do not directly interact, instead H2A.Z recruits the DME homologue ROS1 to its targets in vegetative tissues (Nie et al., 2019). This is reminiscent of the activity of DME in the central cell, where FACT is required for DME access to certain targets. Evidence exists to suggest that DME and FACT interact directly, and so may not require intermediary proteins (Frost et al., 2018). Interestingly, our analysis of promoter DNA methylation at H2A variants showed that the H2A.Z.4 locus may be regulated by DME in the developing seed (Supplementary Figure S1) (Ibarra et al., 2012;Park et al., 2016).

It was previously shown that the *HTA3* and *HTA5* gene promoters exhibited differences in activity, with *HTA5* observed to be less active in the floral bud (Yi et al., 2006). Consistent with this, we show that whilst HTA5 is the predominant protein isoform expressed in the sporophyte, HTA3 predominates in the developing gametophytes, though both are highly expressed in pollen. One explanation for H2A3/5 high expression in the vegetative and central cells is that DME activity creates AP sites during BER, that may lead to the formation of double strand breaks, thereby requiring high levels of H2A.X (Sczepanski et al., 2010). Intriguingly, however, in heterochromatin, the mechanism of DNA repair is different; an H2A.W variant, H2A.W.7 is phosphorylated by ATM to initiate the response in constitutive heterochromatin to DNA damage (Lorkovic et al., 2017).

We observed a significant increase in root hair length in *h2a.x* mutants compared to wild-type in the absence of any DNA damaging conditions. Intriguingly, reduction in H2A.Z incorporation into chromatin also results in an increase in root hair length, since the altered chromatin state mimics phosphate deficiency - activating phosphate deficiency response gene locus (Deal et al., 2005;Smith et al., 2010). In a similar context, *h2a.x* mutations may indirectly affect the expression of root hair-growth genes (Won et al., 2009;Hwang et al., 2017;Mangano et al., 2017). Defective *H2A.X* expression may also cause nutrient-stress, resulting in the modulation of genes involved in root hair growth.

The mechanism of DNA methylation loss in *h2a.x* endosperm remains unclear. Since we identified hypomethylation on both male and female endosperm alleles, but not in embryo, hypomethylation must occur post-fertilization in endosperm, at least on the paternal allele. On the maternal allele, both pre- and post-fertilization DNA methylation dysregulation may be present. Endosperm is a highly unique tissue; it is triploid, and the site of parental competition for generational resources, in part reflected in the activities of DME and FACT in the central cell, which confer deep hypomethylation. Endosperm exhibits highly distinct higher-order chromatin structure compared to other tissues, being less condensed and subsequently featuring increased trans-interactions, increased expression of TEs, and encroachment of heterochromatin into euchromatic regions (Baroux et al., 2007;Yadav et al., 2021). As such, the role of H2A.X in endosperm chromatin may well be distinct from that in other tissues.

Our embryo minus endosperm DMR analyses revealed the following characteristics of *h2a.x*-specific hypomethylated DMRs: enriched in chromatin states 4, 8 and 9 (Sequeira-Mendes et al., 2014) and located in TEs, pericentric heterochromatin and intergenic regions. Chromatin states 4 and 8 are strikingly enriched in intergenic DNA (66.2 and 58.2 %, respectively, (Sequeira-Mendes et al., 2014). Chromatin state 4 is also characterized by the presence of histone variants H3.3 and H2A.Z, and high levels of H3K27me3, but is not highly associated with active transcription. It is also noted to likely contain distal promoters and regulatory elements. Chromatin states 8 and 9 are highly enriched in TEs, and feature H3.1, H3K9me2, and H3K27me1, and although state 8 is a transitional, more decondensed state, they both represent Arabidopsis heterochromatin (Sequeira-Mendes et al., 2014). The enrichment of DNA hypomethylation of these regions in *h2a.x* mutants is intriguing. Whilst *h2a.x* hypomethylated DMRs exhibit overlap with DME target sites, they also represent additional regions not normally demethylated, and their presence in intergenic chromatin and in heterochromatic TEs may be indicative of normal requirement of H2A.X for DME exclusion from these regions; that is, to prevent inappropriate remodeling of regulatory DNA and heterochromatic TEs.

Nucleosome cores are crucial for nuclear DNA organization and function, and a complete loss of *H2A.X* is likely quickly replaced by other H2A variants, such as H2A.Z, H2A.W, or by canonical H2A. Intriguingly, in human cells, H2A.X phosphorylation increases the accessibility of chromatin to DNA methylases, (Heo et al., 2008). We therefore speculate that substitution for other H2A variants in *h2a.x* mutant endosperm contributes to endosperm hypomethylation, perhaps by allowing DME to access loci normally not permitted.

In conclusion, we demonstrate that *H2A.X* is expressed widely in developing *Arabidopsis* tissues and gamete companion cells, and show that the DNA damage response is impaired in *h2a.x* mutant roots and seedlings. We show that *h2a.x* mutant endosperm exhibits DNA hypomethylation at intergenic regions and heterochromatic TEs, creating a large number of embryo-endosperm DMRs, not present in wild-type. We hypothesize that the presence of H2A.X in endosperm contributes to a chromatin structure that is refractive to DME and DME-FACT activity, preventing inappropriate DNA demethylation in intergenic and heterochromatic DNA.

## Data Availability Statement

The datasets for this study can be found in GEO accession (TBA)

## Author Contributions

RLF, YC and JMF conceived the project; JMF, JL, PHH, SJHL, YM, MB, AMR, HTC performed the experiments; JMF, YC, TFH, and RLF analyzed the data; JMF, YC, and RLF wrote the article with contributions of all the authors. All authors contributed to the article and approved the submitted version.

## Acknowledgments

We thank Christina Wistrom and formerly Barbara Rotz for their management of the University of California, Berkeley, Oxford Tract greenhouse facility. We are grateful to Christian Ibarra for his advice on manual endosperm dissection. This work used the Vincent J. Coates Genomics Sequencing Laboratory at the University of California, Berkeley, supported by NIH S10 Instrumentation Grants S10RR029668, S10RR027303, and S10OD018174, and the authors would like to particularly thank Shana McDevitt for her assistance.

## Supplementary Materials

**Supplementary Figure S1. Analysis of CG DNA methylation at H2A variant genomic loci in WT (Col-0) and *dme-2* mutant Arabidopsis**.

(A) *HTA3* (H2A.X). (B) *HTA5* (H2A.X). (C) *HTA4* (H2A.Z). CC = Central cells (Park et al., 2016), Endo = Endosperm (Hsieh et al., 2009). Transcription start site and promoter regions are highlighted with orange box. All cytosine methylations are included without read cutoff. All bars of histogram indicate the methylation % level of single cytosine. Bismark was used for bisulphite sequencing data read alignment and Seqmonk was used for visualization.

**Supplementary Figure S2. Genome-wide non-CG methylation analysis of *h2a.x* mutant developing embryo, endosperm and seedling**.

Ends analysis of selfed *h2a.x* mutant and segregating WT genomic CHG methylation in genes (A) and TEs (B), as well as CHH methylation in genes (C) and TEs (D), with those aligned according to their 5’ and 3’ ends. Data for seedling, endosperm and embryo (linear-bending cotyledon) are shown. Endosperm and embryo are hypomethylated at CHH at gene edges and in TEs. Since embryo CHH methylation levels are incredibly sensitive to gestational age, they may be indicative of slightly later dissection of WT for this sample.

**Supplementary Figure S3. Allele-specific genome-wide CG methylation analysis of *h2a.x* mutant developing embryo and endosperm**.

Female WT Col-0 or *h2a.x* homozygous mutants were crossed with WT Ler pollen and the methylation levels in F1 seeds were analyzed. ‘*h2a.x* paternal’ denotes a WT paternal allele now resident in a heterozygous *h2a.x* mutant seed. Ends analysis of embryo and endosperm CG genomic methylation in genes (A and B, respectively) as well as those in TEs (C and D, respectively) are shown, with genes and TEs aligned according to their 5’ and 3’ ends. The maternal *h2a.x* endosperm allele is hypomethylated compared to WT. Ends analysis of embryo and endosperm CHG genomic methylation in genes (E and F, respectively) as well as those in TEs (G and H, respectively) are shown, with genes and TEs aligned according to their 5’ and 3’ ends. Ends analysis of embryo and endosperm CHH genomic methylation in genes (I and J, respectively) as well as those in TEs (K and L, respectively) are shown, with genes and TEs aligned according to their 5’ and 3’ ends. CHH methylation in genes at intergenic regions is decreased on both *h2a.x* endosperm alleles compared to WT. CHH methylation in TE bodies is decreased on both *h2a.x* embryo and endosperm alleles compared to WT.

