## Supplementary Figures for "*H2A.X* mutants exhibit enhanced DNA demethylation in *Arabidopsis thaliana*"

Frost et al., H2A.X
Supplementary Figures 1-3

Figure S1 – (A) Central cell and endosperm DNA methylation at HTA.X.3 locus

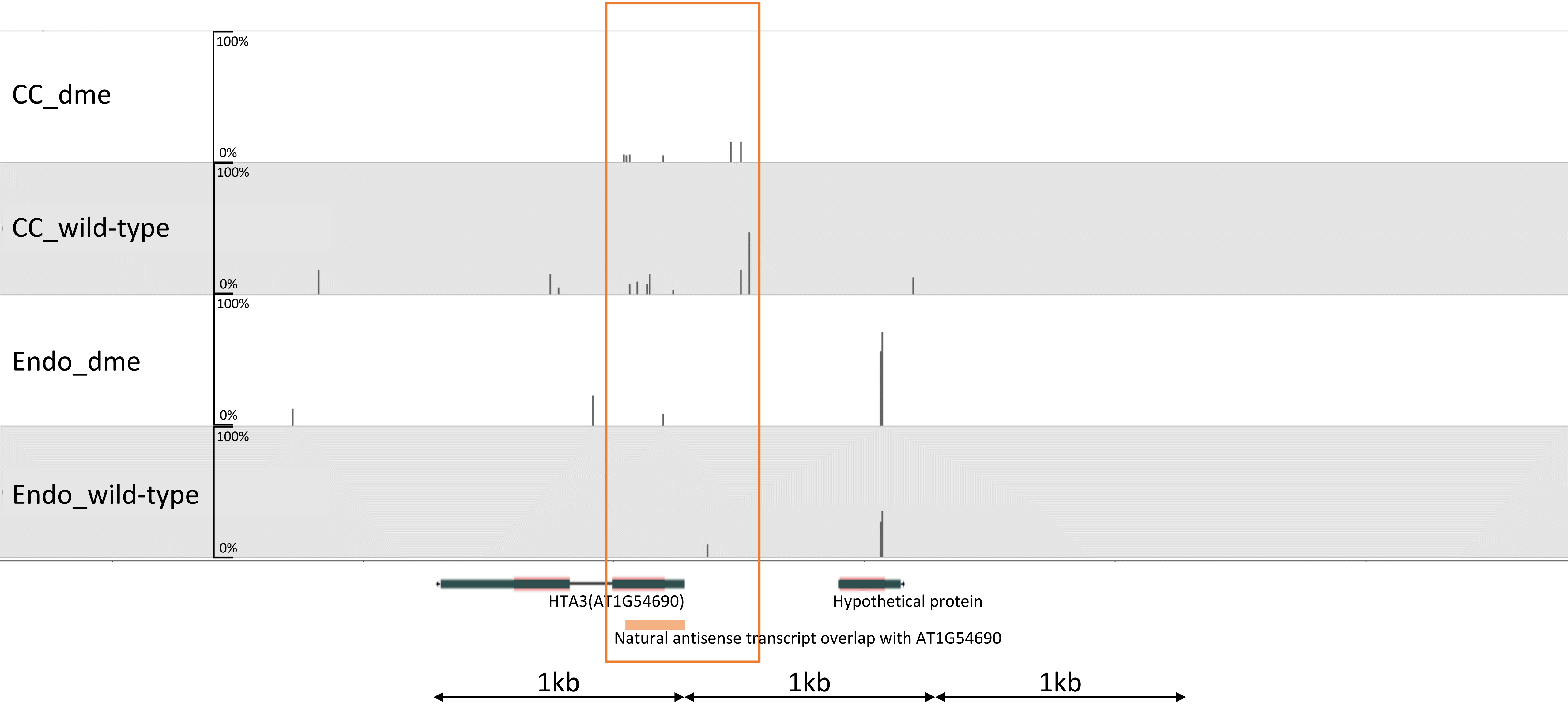

Figure S1 – (B) Central cell and endosperm DNA methylation at HTA.X.5 locus

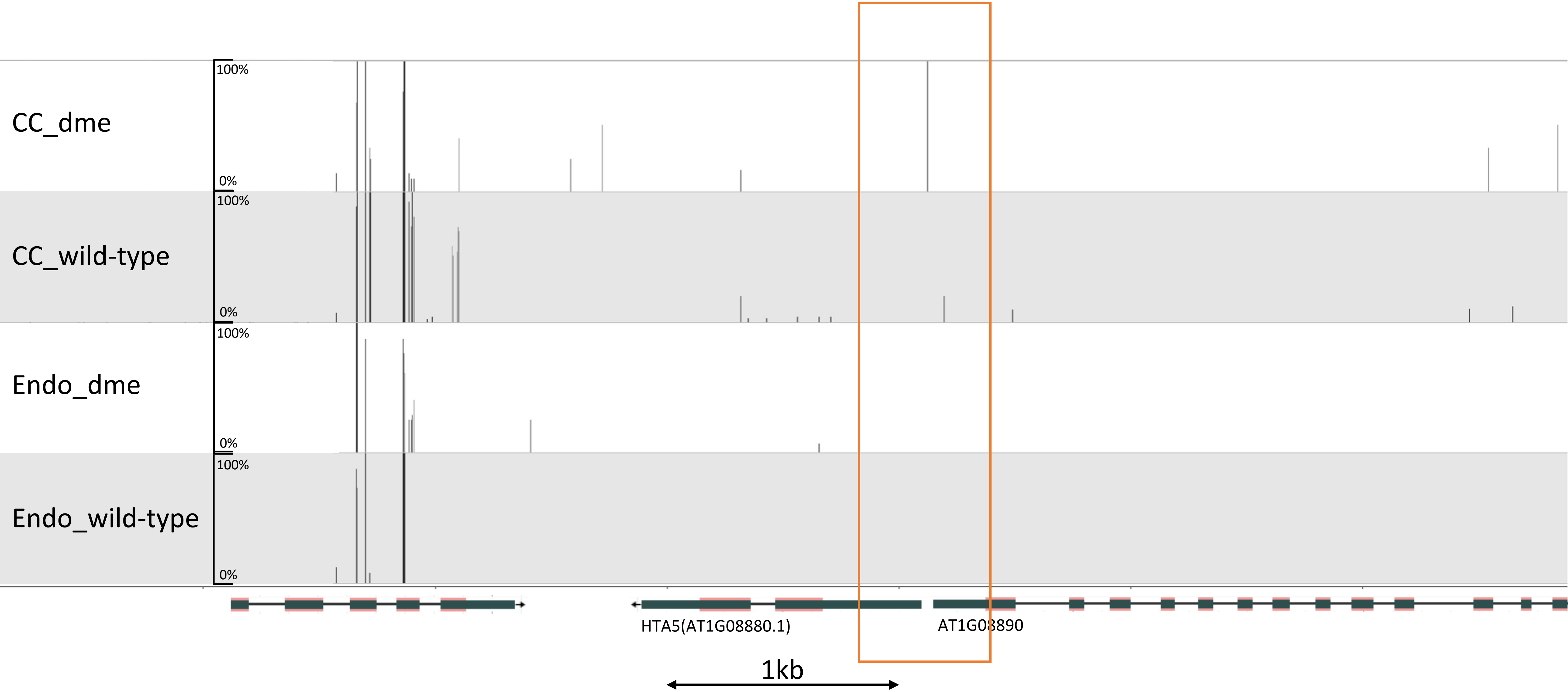

Figure S1 - (C)– H2A.Z.4 – promoter hypermethylation in dme-2 mutant CC and endosperm

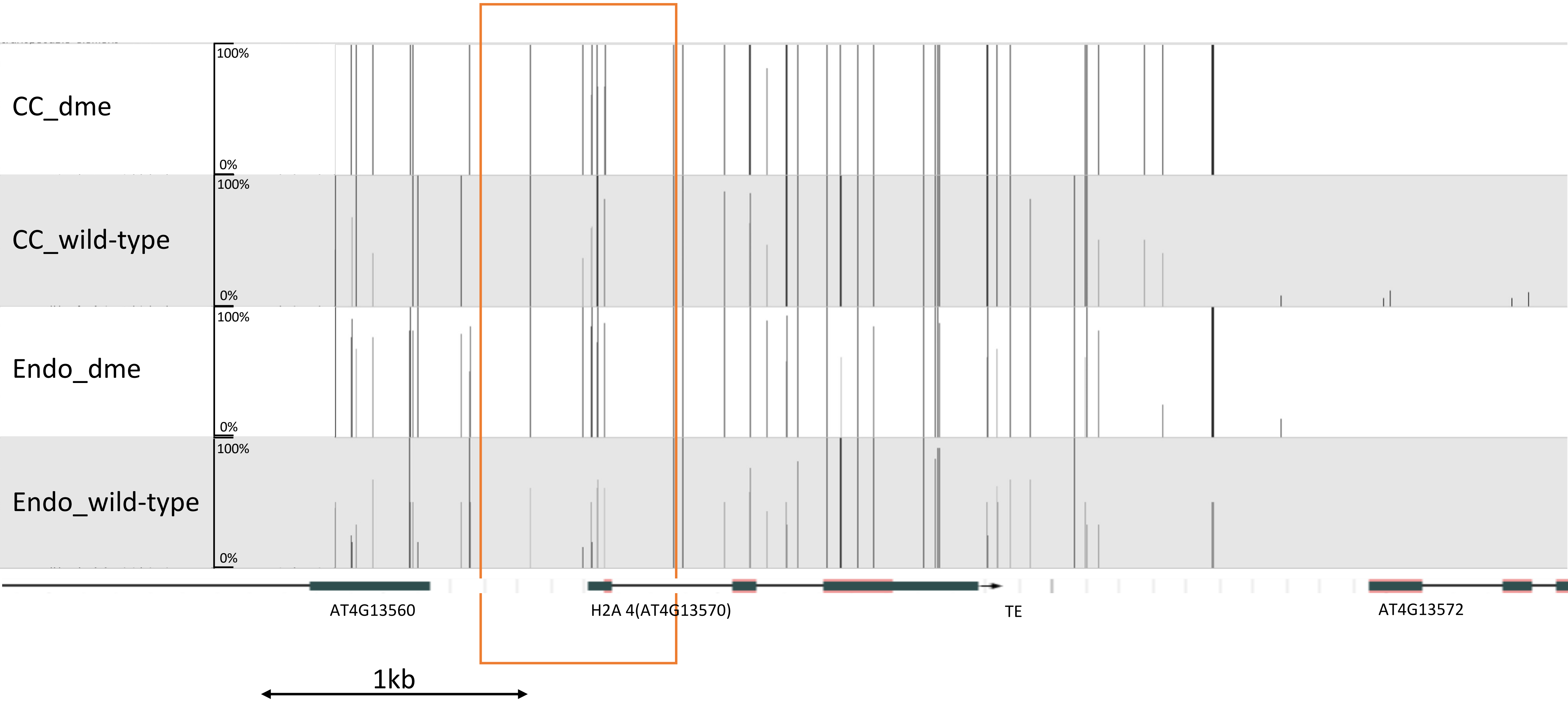

### Figure S2 – Methylome analysis of homozygous H2AX mutant seeds and seedlings, non-CG methylation

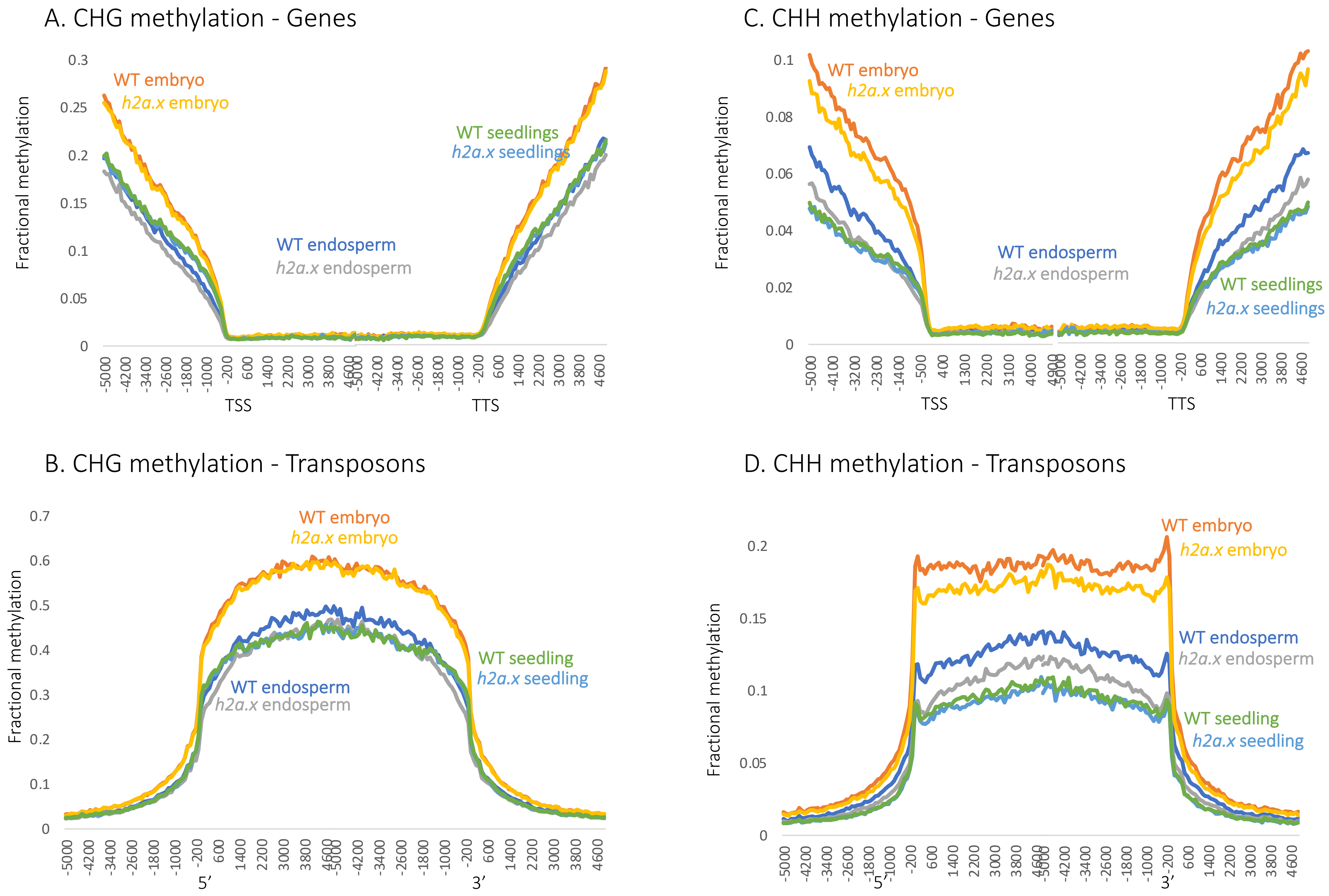

Figure S3 – CG Methylome analysis of H2AX mutants – maternal vs paternal plots

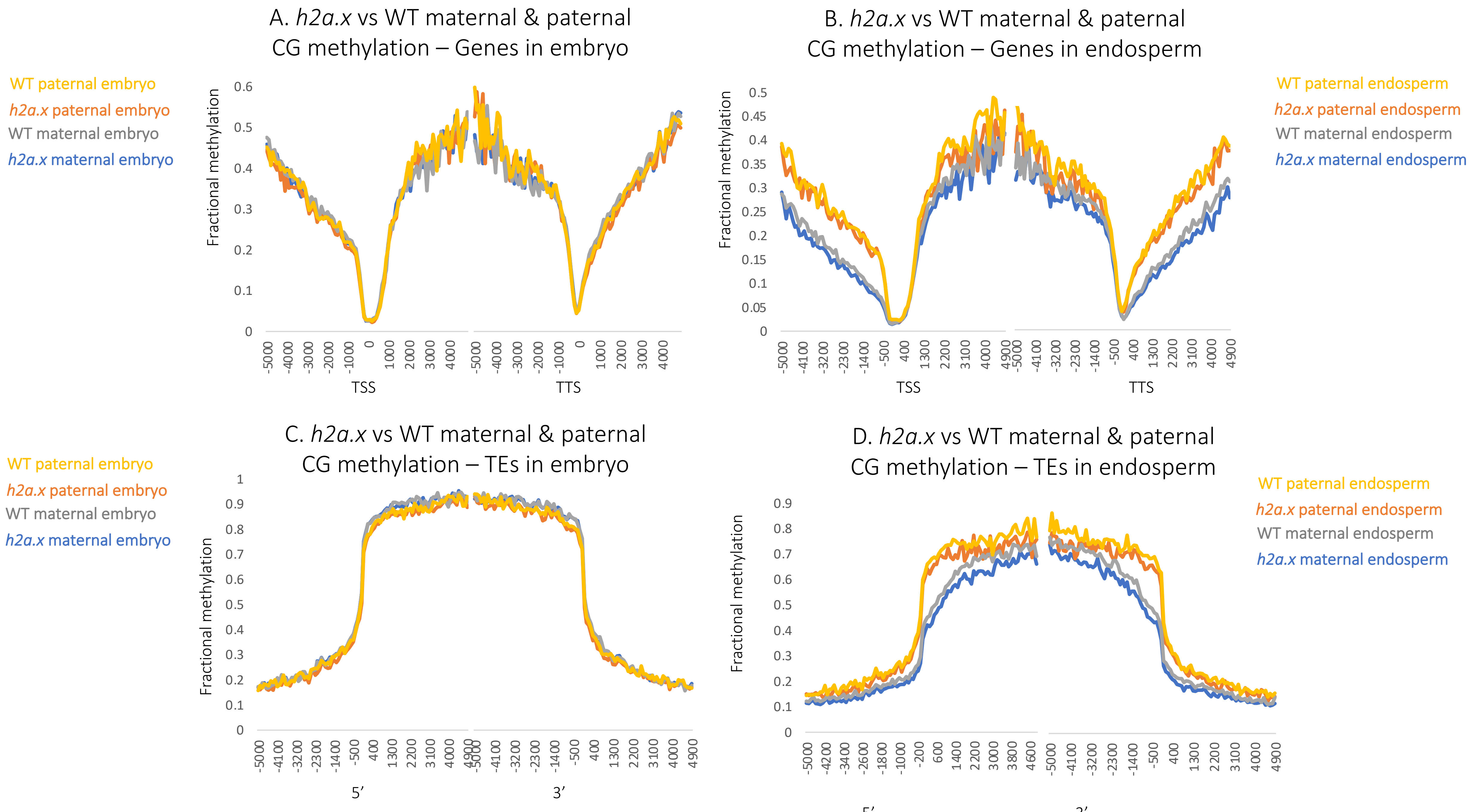

### Figure S3 – CHG Methylome analysis of H2AX mutants – maternal vs paternal plots

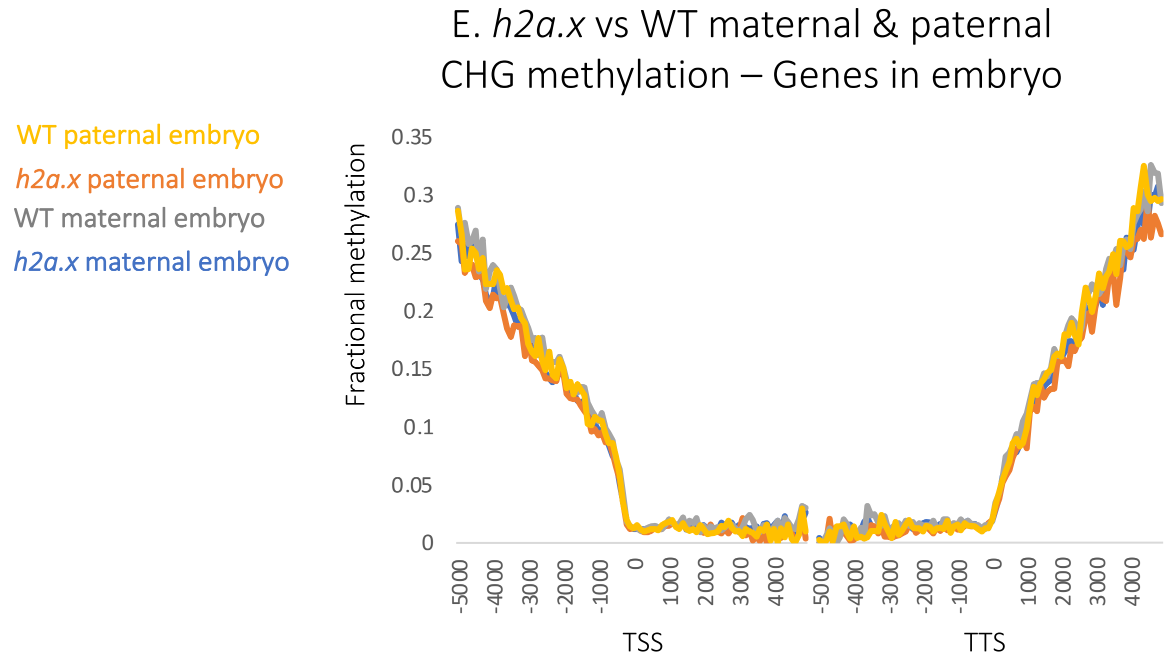


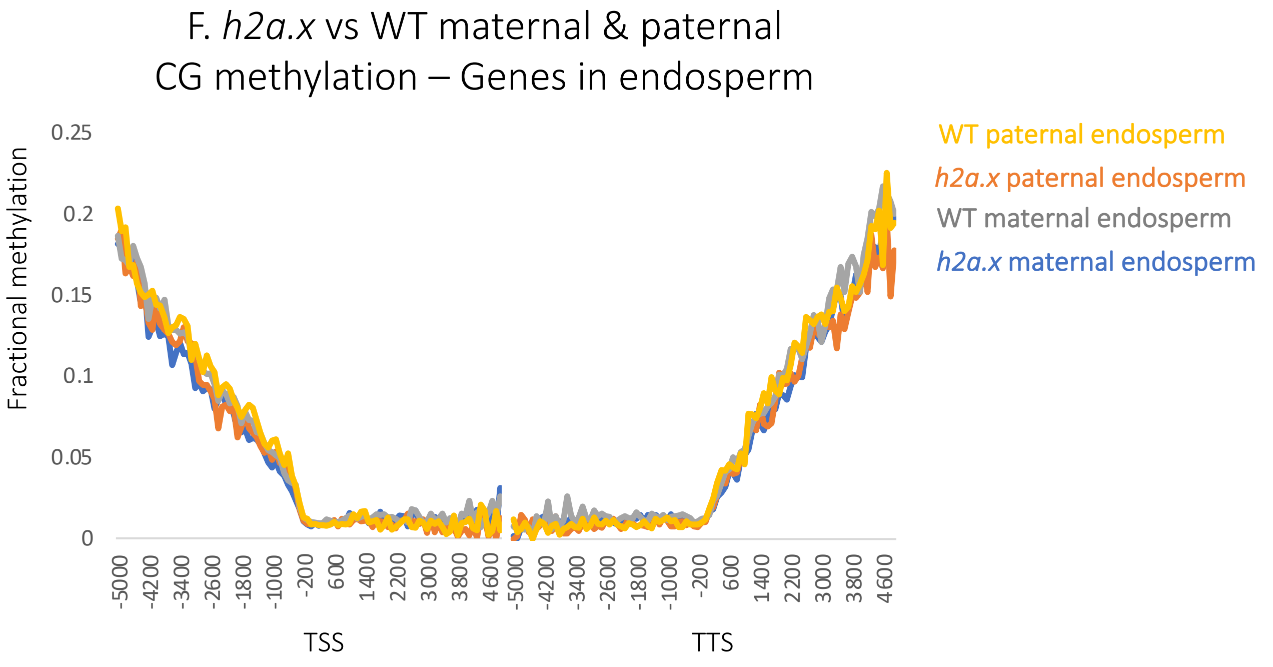


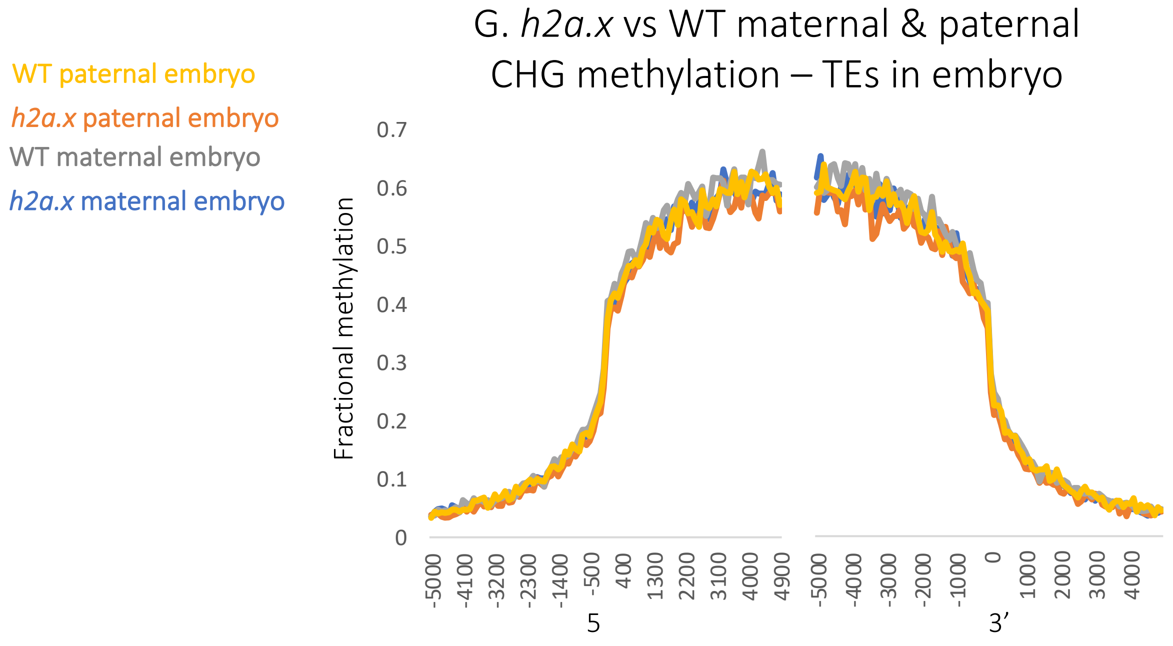


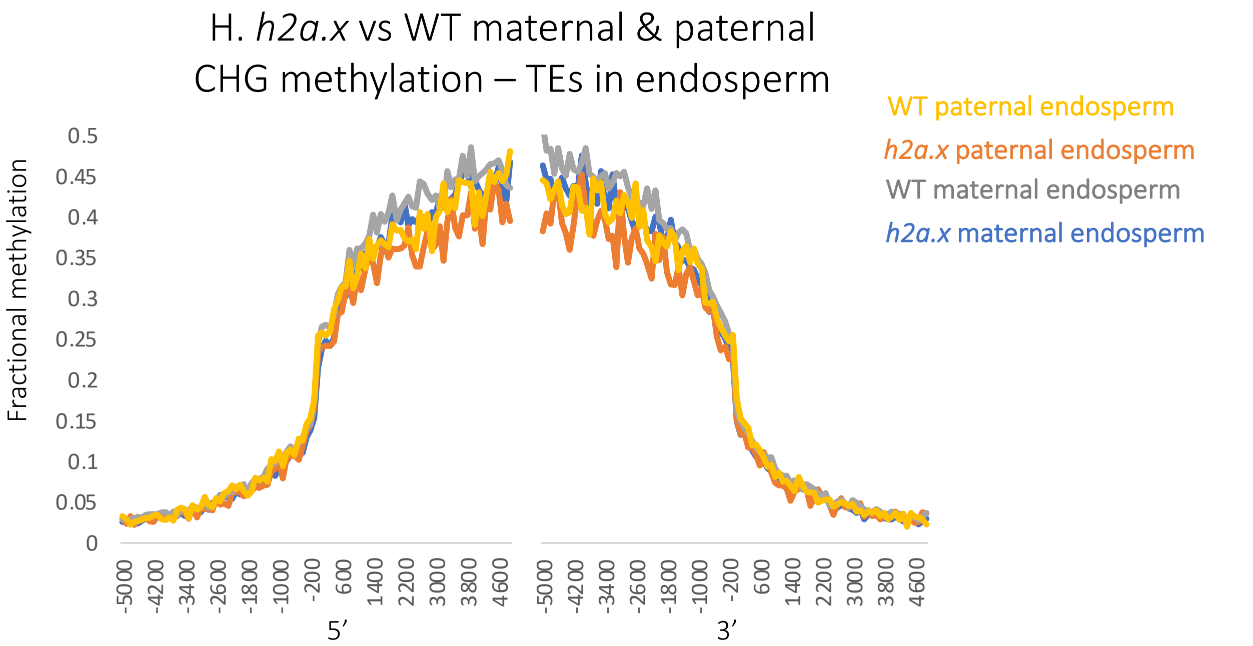

Figure S3 – CHH Methylome analysis of H2AX mutants – maternal vs paternal plots

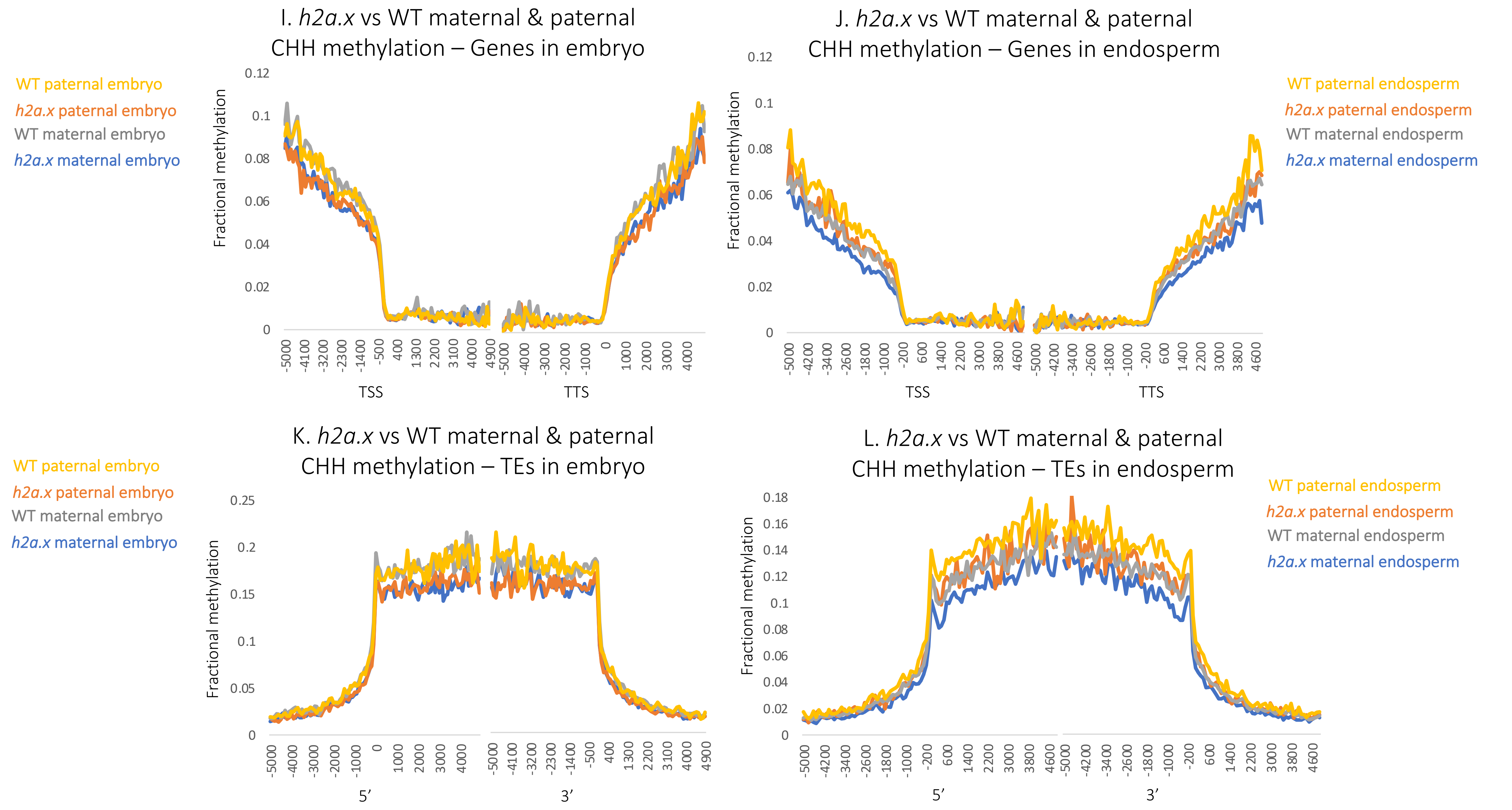
